# Frequency domain analysis of fluctuations of mRNA and protein copy numbers within a cell lineage: theory and experimental validation

**DOI:** 10.1101/2020.09.23.309724

**Authors:** Chen Jia, Ramon Grima

## Abstract

The stochasticity of gene expression is manifested in the fluctuations of mRNA and protein copy numbers within a cell lineage over time. While data of this type can be obtained for many generations, most mathematical models are unsuitable to interpret such data since they assume non-growing cells. Here we develop a theoretical approach that quantitatively links the frequency content of lineage data to subcellular dynamics. We elucidate how the position, height, and width of the peaks in the power spectrum provide a distinctive fingerprint that encodes a wealth of information about mechanisms controlling transcription, translation, replication, degradation, bursting, promoter switching, cell cycle duration, cell division, and gene dosage compensation. Predictions are confirmed by analysis of single-cell *Escherichia coli* data obtained using fluorescence microscopy. Furthermore, by matching the experimental and theoretical power spectra, we infer the temperature-dependent gene expression parameters, without the need of measurements relating fluorescence intensities to molecule numbers.

## Introduction

In recent years, measurements of the size, division events, and the content of single cells over extended time (many generations) has been made possible due to advances in microfluidic devices and live-cell imaging [1–3]. While the existing data are typically for proteins, such measurements are also in principle possible for mRNAs particularly with the advent of new methods to visualize RNA dynamics in live cells using bright and stable fluorescent RNAs [4]. The data in these experiments are sampled at a rate that is much higher than the frequency of cell division thus providing us with a means to understand the temporal variation of gene expression as a cell progresses through its cell cycle.

A common feature of these time traces is a noisy oscillatory variation of the fluorescence (from fluorescently labelled proteins) with time with a period that is roughly coincident with the interval between two successive cell division events. A sawtooth type of temporal pattern is expected due to a sharp dip in the protein numbers at cell division stemming from the partitioning of the contents of the mother cell amongst two daughter cells. While these oscillations are regular in some cases, very often they display a significant degree of noisiness. This reflects the inherent stochasticity of transcription, translation, and replication [5–7], noise introduced or modified by homeostatic mechanisms such as those that compensate for the doubling of gene copies at replication [8, 9], and non-genetic sources of noise such as variability in the duration of the cell cycle from one generation to the next [1, 10, 11] and variability in the number of molecules allocated to a new born cell at cell division [12, 13]. Hence it follows that a measure of the regularity of an oscillation, such as the power spectrum of fluorescence fluctuations calculated over a lineage, encapsulates within it a large amount of information about the inherent chemical and physical processes, both deterministic and stochastic, that shape cellular dynamics.

An essential first step to link the properties of the power spectrum to the underlying dynamic intracellular processes is the derivation of a mathematical formula for the spectrum as a function of gene expression rate parameters. For this purpose, the standard stochastic models in the literature that are based on the two-stage or three stage-models of gene expression [14] are not useful because they do not provide a mathematical description of processes along a cell lineage. These model transcription and translation, and implicitly model dilution due to cell division via an effective decay reaction; however simulated time traces of the protein numbers based on these models using the stochastic simulation algorithm will not display any noisy oscillatory behaviour since the partitioning of molecules at cell division is not taken into account. This also implies a lack of description of important cell cycle features such as the high variability in the interdivision time that is a characteristic of experimental time traces. Recently models that surmount the aforementioned limitations of the standard models have been studied leading to mathematical formulae for the mean and variance of mRNA/protein numbers [15–17] and also for the steady-state distributions of these numbers [18, 19] calculated across a cell lineage. These statistical measures provide different information than the power spectrum; notably the former, unlike the latter, neither provide an understanding of the correlations between molecule numbers at two time points nor of the frequency composition of fluctuations in molecule numbers [20].

In this article, for the first time, we calculate in closed-form the power spectrum of fluctuations across a lineage in a stochastic gene expression model with a high level of biological realism, including a description of transcription, translation, degradation, bursting, promoter switching, DNA replication, gene dosage compensation, and symmetric/asymmetric partitioning at cell division. The analytical expressions give insight into how the regularity and noisiness of the oscillations in the mRNA/protein abundance across generations are related to the rate parameters associated with the various subcellular processes at play; the theory also makes various predictions that are then verified by analysis of a publicly available single-cell data set of *Escherichia coli* followed over 70 generations for three different growth conditions; finally we show how matching the experimental and theoretical power spectra enables an accurate inference of all the rate parameters.

## Results

### Model specification

We consider a detailed model of stochastic gene expression in an individual cell which includes promoter switching, transcriptional or translational bursting, cell cycle duration variability, gene replication, gene dosage compensation, and symmetric or asymmetric cell division (see Fig. 1(a) for an illustration). The specific meaning of all model parameters is listed in Table 1 and the biological values of some key parameters in different cell types are listed in table S1. The model is based on a number of assumptions that are closely tied to experimental data. The assumptions are as follows.

**Fig. 1.**
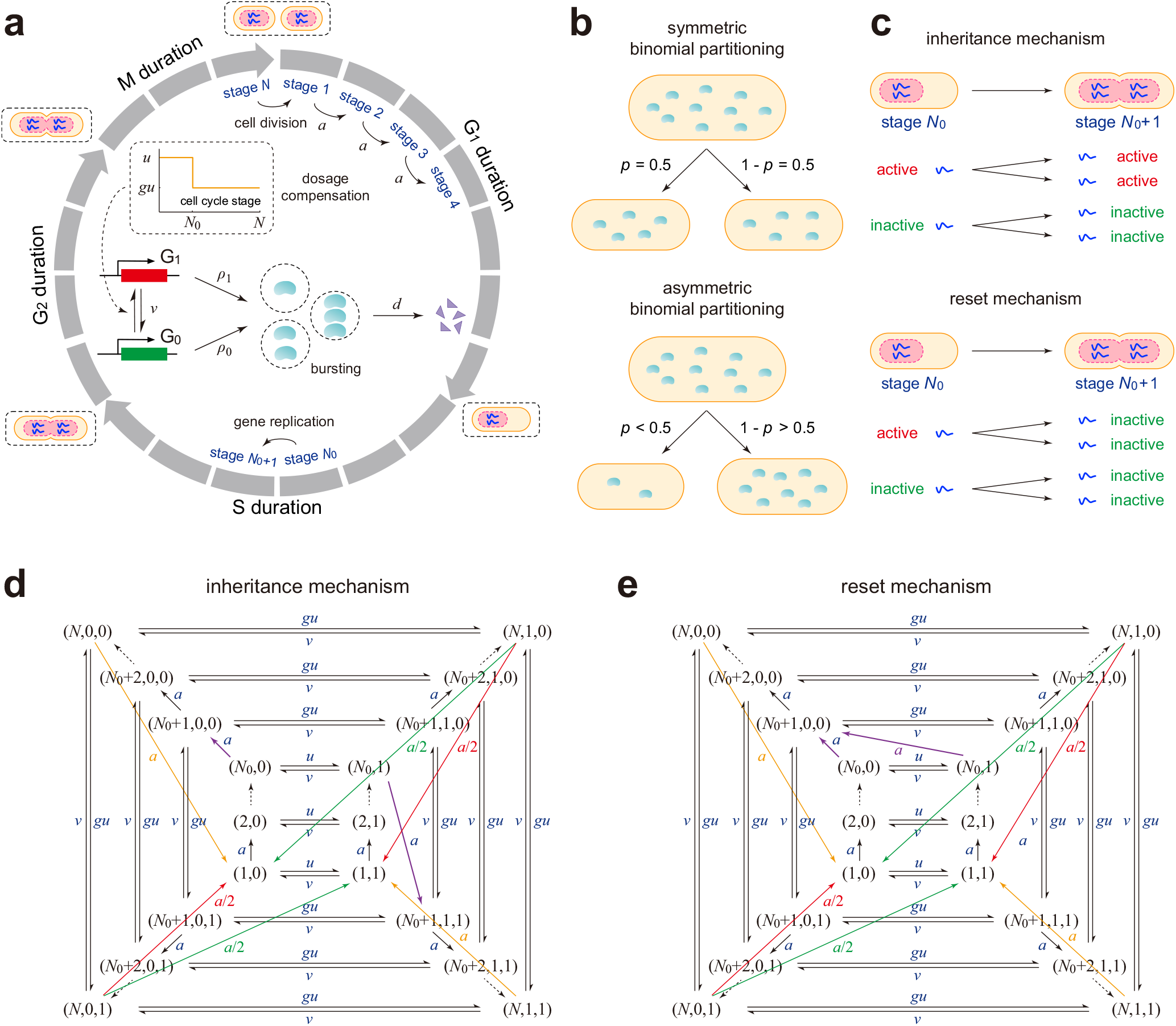
The model. (**a**) Schematic illustrating the model describing *N* effective cell cycle stages, gene replication at stage *N*_0_, promoter switching between active (red) and inactive (green) states, bursty production of the gene product in the two gene states, degradation, and gene dosage compensation induced by a decrease in the activation rate of the gene after replication (see inset graph). (**b**) At stage *N*, a mother cell divides into two daughters that are typically different in size (asymmetric division) with the larger daughter inheriting more molecules. Symmetric division is the special case where the daughters are equisized. (**c**) At replication, the gene states of the two daughter copies can be the same as that of the mother copy (inheritance mechanism) or else they are both reset to the inactive state (reset mechanism). (**d**),(**e**) Transition diagram of cellular states under the two mechanisms. Before replication, a cellular state can be represented by an ordered pair (*r, i*), where *r* is the cell cycle stage and *i* is the gene state of the mother copy; after replication, a cellular state can be represented by an ordered triple (*r, i, j*), where *i* and *j* are the gene states of the two daughter copies. See main text for explanation of the colored arrows.

**Table 1.** Model parameters and their meaning.

| Meaning of model parameters |  |
| --- | --- |
| $u$ | gene switching rate from OFF to ON before replication |
| $gu$ | gene switching rate from OFF to ON after replication |
| $v$ | gene switching rate from ON to OFF |
| $\rho_0$ | burst production rate when the gene is OFF |
| $\rho_1$ | burst production rate when the gene is ON |
| $B$ | mean burst size of the gene product |
| $d$ | degradation rate of the gene product |
| $N$ | number of cell cycle stages |
| $N_0$ | number of cell cycle stages before replication |
| $w$ | proportion of the cell cycle before replication |
| $a$ | transition rate from one cell cycle stage to the next |
| $f$ | cell cycle frequency |
| $\eta$ | ratio of the degradation rate and the cell cycle frequency |
| $p$ | allocation probability in asymmetric binomial partitioning |
| $\rho_{\text{eff}}$ | effective burst production rate before replication |
| $\kappa\rho_{\text{eff}}$ | effective burst production rate after replication |
| $d_{\text{eff}}$ | effective decay rate of the gene product |

1) The promoter of the gene of interest can switch between an inactive state *G*_0_ and an active state *G*_1_ with switching rates *u* and *v* before gene replication [14]. Dosage compensation is modeled as a change in the switching rate of the promoter from the inactive to the active state upon replication with its value being *u* before replication and *gu* after replication. This assumption is supported by experiments [8].

2) In each gene state *G*_*i*_ (*i* = 0, 1), the synthesis of the gene product of interest, mRNA or protein, occurs at a rate *ρ*_*i*_ in bursts of a random size sampled from an arbitrary probability distribution *µ* = (*µ*_*n*_). This means that in each burst, there is a probability *µ*_*n*_ of producing *n* copies of the gene product. In previous papers, the synthesis of mRNA *in each gene state* is assumed to be non-bursty [21], i.e. the mRNA molecules are produced one at a time. In this case, *µ*_*n*_ = *δ*_1,*n*_ is the Kronecker delta which takes the value of 1 when *n* = 1 and the value of 0 otherwise. On the other hand, in models where mRNA is not explicitly described, the effective synthesis of protein in each gene state is usually assumed to be bursty with the burst size sampled from a geometric distribution 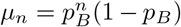, where *p*_*B*_ = *B/*(1 + *B*) with *B* being the mean burst size; this is due to rapid synthesis of protein molecules from a single short-lived mRNA molecule [22, 23]. Therefore, the arbitrariness of the burst size distribution allows us to analyze the dynamics of both mRNA and protein in a unified model.

3) The gene product is degraded via first-order kinetics with rate constant *d*, which is a common assumption supported by experiments [24].

4) Each cell can exist in *N* effective cell cycle stages, denoted by 1, 2, *…, N*, with *a* being the transition rate from one stage to the next, which is assumed to be the same for all stages. Since the transition time between stages is exponentially distributed, the duration of the cell cycle is Erlang distributed with mean *N/a* and thus the cell cycle frequency is *f* = *a/N*. In our model, the noise in the doubling time, characterized by the coefficient of variation squared, is equal to 1*/N*. As *N* → ∞, the noise vanishes and thus the doubling time becomes fixed. Hence, our model allows the investigation of the influence of cell cycle duration variability on stochastic gene expression.

We emphasize that the effective cell cycle stages introduced here do not directly correspond to the four biological cell cycle phases of eukaryotic cells (G_1_, S, G_2_, and M) since the durations of the latter are typically not exponentially distributed. In our model, a cell cycle phase corresponds to multiple effective cell cycle stages (Fig. 1(a)). By introducing a number of effective cell cycle stages, our model has the property that the total duration of the cell cycle and the durations of individual cell cycle phases are all Erlang distributed. This is in agreement with experiments in various cell types [17, 25–28].

5) Cell division occurs when the cell transitions from effective stage *N* to the next stage 1. At division, most previous papers assume that the mother cell divides into two via symmetric binomial partitioning: each molecule has an equal chance to be allocated to one of the two daughters [19, 29]. However, asymmetric cell division is common in biology [13, 30]. For instance, *Saccharomyces cerevisiae* divides asymmetrically into two daughters with different sizes. *Escherichia coli* may also undergo asymmetric division with old daughters receiving fewer gene products than new daughters [31]. Here we extend previous models by considering asymmetric binomial partitioning at cell division: the probability for a molecule being allocated to one daughter is *p* ≤ 1*/*2 and the probability of being allocated to the other is *q* = 1 − *p* (Fig. 1(b)). After cell division, we randomly track one of the two daughters with probability 1*/*2; hence our model corresponds to cell lineage measurements performed using a mother machine such as in [32].

6) The replication of the locus containing the gene of interest occurs over a period that is much shorter than the rest of the cell cycle. Note that the replication of the whole genome within a cell cannot be assumed to be instantaneous. However, since the replication time of a particular gene is much shorter than the total duration of the S phase, it is reasonable to consider it to be instantaneous. Specifically, we assume that gene replication occurs instantaneously when the cell transitions from a fixed effective stage *N*_0_ ∈ [1, *N* − 1] to the next stage *N*_0_ + 1. We shall refer to the gene copy that is replicated as the mother copy and to the duplicated gene copies as the daughter copies. Under this assumption, for haploid cells, there is only one mother copy during the first *N*_0_ stages and two daughter copies during the last *N* − *N*_0_ stages; for diploid cells, the number of gene copies varies from two to four upon replication. For diploid cells, we assume that the two alleles act independently of each other [33, 34].

7) After replication, the two daughter copies can either inherit the gene state from the mother copy or be both reset to the inactive state [18]. To distinguish between them, we refer to the former as the inheritance mechanism and the latter as the reset mechanism (Fig. 1(c)). The consideration behind the former is the copying of the landscape of histone modifications (implicated in gene activation) during DNA replication [35]. One plausible explanation for the latter is that to avoid the potential risk of conflict between replication and transcription [36], it is likely that in the region where replication is ongoing or just completed, there is no transcription, indicating an inactive state.

We next describe our stochastic model for haploid cells. The results for diploid cells can be easily deduced from the haploid case using allelic expression independence (see Methods). Based on the stage of cell cycle progression and the states of the gene copies, there are many possible cellular states for a cell. Before replication, the cell can exist in 2*N*_0_ cellular states that can be represented by an ordered pair *α* = (*r, i*), where *r* ∈ [1, *N*_0_] is the cell cycle stage and *i* = 0, 1 is the gene state of the mother copy; after replication, the cell can exist in 4(*N* − *N*_0_) cellular states that can be represented by an ordered triple *α* = (*r, i, j*), where *r* ∈ [*N*_0_ + 1, *N*] is the cell cycle stage and *i, j* = 0, 1 are the gene states of the two daughter copies. In sum, there are a total of *K* = 4*N* − 2*N*_0_ possible cellular states.

The evolution of cellular state transitions is governed by a Markovian model whose master equation is given in section S1. The transition diagrams of the Markovian model under the inheritance and reset mechanisms are illustrated in Fig. 1(d),(e), respectively. The purple arrows show that upon replication, the cellular state will transition from (*N*_0_, *i*) to (*N*_0_ + 1, *i, i*) for the inheritance mechanism and from (*N*_0_, *i*) to (*N*_0_ + 1, 0, 0) for the reset mechanism. After division, we randomly track one of the two daughters and thus the cell will transition from (*N, i, j*) to (1, *i*) or (1, *j*). The orange arrows illustrate those transitions from (*N, i, i*) to (1, *i*) with rate *a*, the red arrows illustrate those from (*N, i, j*), *i ≠ j* to (1, *i*) with rate *a/*2, and the green arrows illustrate those from (*N, i, j*), *i* ≠ *j* to (1, *j*) with rate *a/*2.

It then follows that the microstate of the gene of interest can be represented by (*α, n*), where *α* is the cellular state and *n* is the copy number of the gene product. The evolution of the complete stochastic gene expression dynamics is governed by a master equation which is given in Methods.

### General expressions for the power spectrum of single-cell measurements across generations

Experiments suggest that the periodicity of the cell cycle can induce oscillatory behavior in gene expression [1]. In fact, if we use a deterministic model to describe the synthesis and degradation of the gene product and assume that the molecule numbers halve after a deterministic doubling time, then the solution of the deterministic rate equation (the time series of gene product abundances) will be periodic under all choices of rate parameters. However, numerous time lapse experiments [6, 37] have shown that the time course data of expression levels in single cells do not always appear oscillatory, due to various sources of noise such as cell cycle duration variability, gene copy number variability, cell division asymmetry, gene expression bursting, and promoter switching. Here we examine how these sources of noise influence the robustness of sustained oscillations.

Let *n*(*t*) denote the copy number of the gene product in a cell at time *t*. Stochastic gene expression oscillations are often characterized by two functions: the autocorrelation function and the power spectrum. The former *R*(*t*) = Cov_ss_(*n*(0), *n*(*t*)) is defined as the steady-state covariance of *n*(0) and *n*(*t*), while the latter 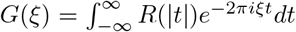 is defined as the Fourier transform of the former, where *ξ* ≥ 0 denotes the frequency. In general, sustained oscillations cannot be observed if the power spectrum *G*(*ξ*) is a monotonically decreasing function of *ξ*. In contrast, a non-monotonic power spectrum with one or more off-zero peaks implies the presence of sustained oscillations; the dominant frequencies of these oscillations are the values of *ξ* at which the modes of the power spectrum occur.

While our model is complex due to the large variety of biological processes that it captures, its autocorrelation function can still be computed exactly as

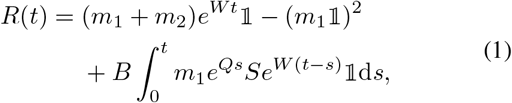

where *m*_1_ = (*m*_1*α*_) and *m*_2_ = (*m*_2*α*_) are two row vectors whose components are the first and second factorial moments of the gene product abundance in all cellular states, 𝟙 is the column vector whose components are all 1, *S* = diag(*ρ*_*α*_) is the diagonal matrix whose diagonal elements are the burst production rates in all cellular states, *Q* = (*q*_*αβ*_) is generator matrix of cellular state transitions (see Fig. 1(d),(e) for the transition diagram which has some orange, red, and green arrows), and *W* = (*w*_*αβ*_) is another matrix obtained from *Q* by replacing the rates of orange arrows by *a/*2, replacing the rates of red arrows by *pa/*2, replacing the rates of green arrows by *qa/*2, and subtracting *d* from the diagonal entries (see Methods for the proof and the detailed expressions of each term).

Note that the autocorrelation function *R*(*t*) in Eq. (1) is expressed in matrix form. A more explicit expression can be obtained by expanding the matrix exponentials *e*^*W t*^ and *e*^*Qs*^ in terms of their eigenvalues and eigenvectors. We find that the autocorrelation function (power spectrum) can be rewritten as the linear combination of 2*K* − 1 exponential (Lorentzian) functions:

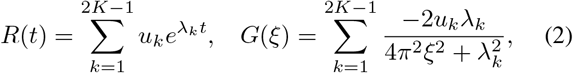

where *K* is the number of cellular states, *λ*_*k*_ (0 ≤ *k* ≤ 2*K* − 1) are all the eigenvalues of the two matrices *Q* and *W*, and *u*_*k*_ are suitable constants (see Methods for the proof and the specific expressions of *u*_*k*_). Since *Q* and *W* are both *K* × *K* matrices, they have a total of 2*K* eigenvalues.

An important special case occurs when promoter switching is much faster than cell cycle progression and gene product degradation, i.e. *f, d* ≪ *u, v* [34]. In this case, gene switching dynamics will reach rapid equilibrium. As a result, the effective burst production rate is given by *ρ*_eff_ = (*ρ*_1_*u* + *ρ*_0_*v*)*/*(*u* + *v*) before replication and given by *κρ*_eff_ = 2(*ρ*_1_*gu* + *ρ*_0_*v*)*/*(*gu* + *v*) after replication, where *κ* ≥ 1 is a factor characterizing the change in the burst production rate due to gene replication and dosage compensation. In the absence of dosage compensation, we have *gu* = *u* and thus *κ* = 2. In the fast switching regime, the two gene states can be combined into a single one and hence the cellular state is only determined by the *N* effective cell cycle stages. In this case, we do not need to distinguish between the inheritance and the reset mechanisms because they lead to the same oscillatory behavior.

We next examine the properties of stochastic oscillations under fast promoter switching. In this regime, *Q* and *W* reduce to *N* × *N* matrices and the power spectrum in Eq. (2) simplifies because all the eigenvalues can be computed explicitly. The eigenvalues of *Q* are given by *λ*_*k*_ = −*a* + *aω*_*k*_ (0 ≤ *k* ≤ *N* − 1) and the eigenvalues of *W* are given by *λ*_*N*+*k*_ = −*d* − *a* + 2^*−*1*/N*^ *aω*_*k*_ (0 ≤ *k* ≤ *N* − 1), where *ω*_*k*_ = *e*^2*πki/N*^ are all the *N* th roots of unity. In addition, the steady-state mean of gene product abundances can be also calculated explicitly as (see section S3 for the proof)

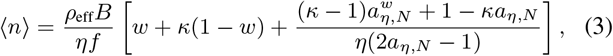

where *w* = *N*_0_*/N* is the proportion of the cell cycle before gene replication, *η* = *d/f* is the ratio of the degradation rate and the cell cycle frequency which serves as a measure for the stability of the gene product, and *a*_*η,N*_ = (1 + *η/N*)^*N*^ ≈ *e*^*η*^ when *N ≫* 1.

To validate the analytical expression of the power spectrum, we compare it with the numerical spectrum calculated by means of the Wiener-Khinchin theorem, which states that 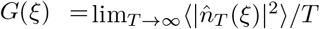, where 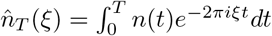 is the truncated Fourier transform of a single stochastic trajectory over the interval [0, *T*] and the angled brackets denote the ensemble average over trajectories (Fig. 2(a)). To guarantee the accuracy of the numerical spectrum, we calculate the time-ensemble average over 5000 stochastic trajectories simulated using Gillespie’s algorithm with the maximum simulation time being chosen as 30*N/a* (about 30 cell cycles). Here we normalize the power spectrum such that *G*(0) = 1. Clearly, the analytical (blue curve) and numerical (red circles) spectra coincide perfectly with each other. In general, the numerical simulations of the power spectrum turn out to be very slow. The analytical solution is hence crucial because it allows a fast exploration of large swathes of parameter space.

**Fig. 2.**
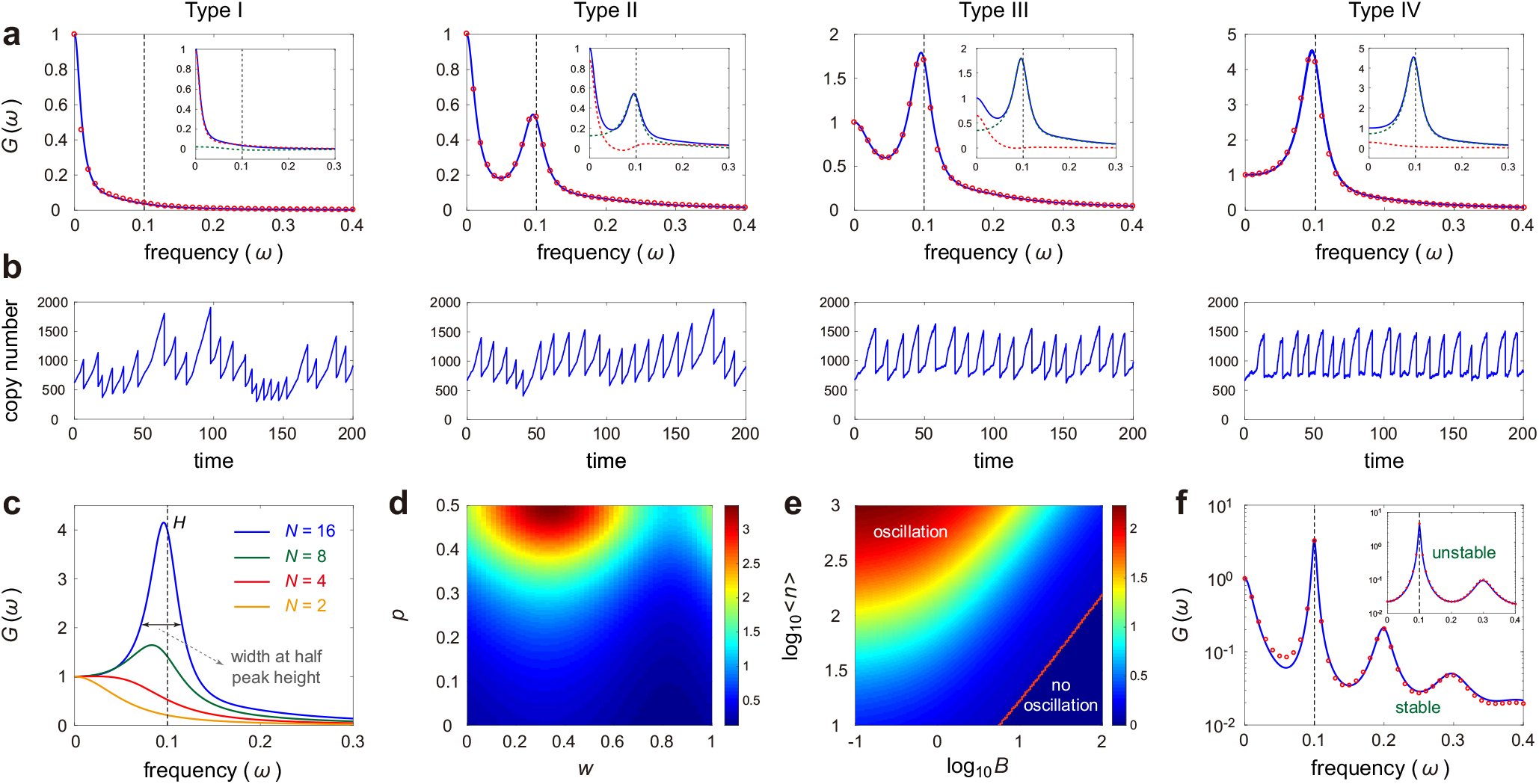
Properties of the power spectrum. (**a**) For significant cell cycle duration variability, the spectra can be of four types. Note that the spectra are normalized such that *G*(0) = 1. Theory (blue) matches stochastic simulations using Gillespie’s algorithm (red circles). Vertical dashed lines show the cell cycle frequency. Insets show the decomposition of the spectrum into two parts controlling the zero and off-zero peaks (red and green dashed curves, respectively). (**b**) Typical trajectories for each type of spectra. (**c**) The off-zero peak becomes higher as cell cycle duration variability decreases (*N* increases). (**d**),(**e**) Heat maps showing the dependence of the height of the off-zero peak (relative to the zero peak) on the parameters controlling the asymmetry of partitioning (*p*), the replication point (*w*), the burstiness of gene expression (*B*), and the mean number of gene product molecules (⟨*n*⟩). Type I spectra are observed in the region marked “no oscillation”. (**f**) For low cell cycle duration variability, higher-order harmonics of the cell cycle frequency appear; for midway replication, an unstable gene product, e.g. most mRNAs, has a spectrum with no peaks at even harmonics, while a stable gene product, e.g. most proteins, has peaks at all harmonics. Note that the blue curves are computed using the analytical expression given in Eq. (2) and the red circles are computed using the approximate expressions given in Eq. (5) for stable products and Eq. (15) for unstable products. See section S10 for the technical details of this figure.

### Single-cell time traces can be classified into four different types according to their power spectra shape

Our gene expression model displays four different types of power spectra (Fig. 2(a)): (i) the spectrum is unimodal and monotonically decreasing with a peak at zero, (ii) the spectrum is bimodal with the height of the off-zero peak less than 1, (iii) the spectrum is bimodal with the height of the off-zero peak greater than 1, and (iv) the spectrum is unimodal and bell-shaped with the height of the off-zero peak greater than 1. For convenience, we refer to (i)-(iv) as type I, II, III, and IV spectra, respectively. The robustness of oscillations increases as the spectrum changes from type I to type IV (Fig. 2(b)); this is since the increasing height of the off-zero peak relative to the zero peak implies increasing power in a narrow range of frequencies centered about the cell cycle frequency. To better understand the analytical solution of the power spectrum, we decompose it into two parts:

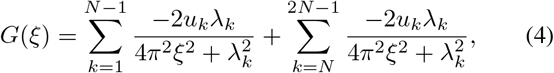

where the first part is the contribution of the eigenvalues of *Q* and the second part is the contribution of the eigenvalues of *W* (insets of Fig. 2(a)). Clearly, the first part (green curves) mainly controls the off-zero peak and the second part (red curves) mainly controls the zero peak as well as the decay of the spectrum. When cell cycle duration variability is small (*N ≫* 1), the first eigenvalue of *W* is given by *λ*_*N*_ = − *d* − *a*(1 − 2^*−*1*/N*^) = − *d* − *a*(1 − *e*^*−*(ln 2)*/N*^) ≈ − *d* − (ln 2)*a/N* = −*d*_eff_, where *d*_eff_ = *d*+(ln 2)*f* is the effective decay rate of the gene product, which is the sum of the decay rate due to active degradation and the decay rate due to dilution at cell division [38]. This explains why the second part characterizes the decay of the spectrum.

### General scaling properties of the height and width of the off-zero spectral peak

To see the effect of cell cycle duration variability on sustained oscillations, we illustrate how the power spectrum given by Eq. (2) varies with *N* (Fig. 2(c)). When *N* is very small, the spectrum only has a peak at zero, implying that no regular oscillations can be observed. However, as *N* increases, the spectrum becomes non-monotonic with the off-zero peak becoming higher and closer to (but still less than) the cell cycle frequency *f* (shown as a vertical dashed line). This indicates that there is a threshold cell cycle duration variability below which the periodicity of the cell cycle leads to sustained oscillations in gene expression.

Moreover, as *f* increases while keeping *N* and *n* fixed, the power spectrum becomes much wider but the height of the off-zero peak remains exactly the same (fig. S1(a)). In fact, if the cell cycle frequency is increased from *f* to *αf* with some *α >* 1, then the coefficients *u*_*k*_ in Eq. (2) will remain the same, but the eigenvalues *λ*_*k*_ will be replaced by *αλ*_*k*_ (see section S2 for the proof). Hence, the power spectrum (autocorrelation function) will be stretched (compressed) along the horizontal axis by a factor of *α*. This explains why the height of the off-zero spectral peak is independent of the cell cycle frequency.

### Symmetric division, midway replication, and non-bursty expression enhance the regularity of oscillations

Oscillations are also affected by the gene replication time, asymmetric cell division, gene expression bursting, and gene expression mean. Fig. 2(d),(e) illustrate the height of the off-zero spectral peak as a function of *w, p, B*, and ⟨*n*⟩. It is clear that the off-zero peak becomes lower as *B* increases and as *p* and ⟨*n*⟩ decrease. The decline of the peak height with increasing *B*, decreasing *p*, and decreasing ⟨*n*⟩ is expected since all of them correspond to an increase in the fluctuations of gene product abundances which counteracts the regularity of oscillations; indeed noise above a certain threshold can even completely destroy sustained oscillations (Fig. 2(e)). This is in sharp contrast to a negative feedback genetic loop, where random bursting can promote the regularity of oscillations [39, 40]. In addition, we also find that sustained oscillations are the most regular when *w* is neither too small nor too large (Fig. 2(d)).

### Contrasting the properties of the off-zero peak for stable and unstable gene products

To further understand these observations, we consider two important special cases. In bacteria and yeast, most proteins have very long half-lives, i.e. *η* ≪ 1 (table S1). For such stable gene products, when cell cycle duration variability is not too large, the power spectrum can be simplified as

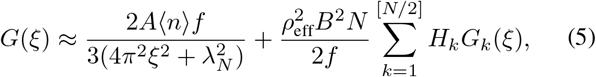

where *A* and *H*_*k*_, *k* ≥ 1 are suitable constants and

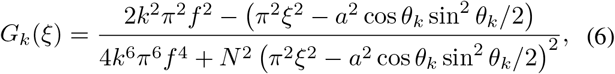

with *θ*_*k*_ = 2*kπ/N* (see Methods for the expressions of *A* and *H*_*k*_ and section S4 for the proof). In Eq. (5), the decay of the power spectrum is mainly controlled by the first term, while the off-zero peak is controlled by the function *G*_1_(*ξ*) in the second term. The influence of the remaining functions *G*_*k*_(*ξ*), *k* ≥ 2 in the second term will be discussed later.

From Eq. (6) with *k* = 1, the position of the off-zero peak is given by *ξ* = (*a/π*) cos *θ*_1_ sin(*θ*_1_*/*2) *< aθ*_1_*/*2*π* = *a/N*, which is smaller than the cell cycle frequency *f* = *a/N*. When *N ≫* 1, the peak position is approximately equal to *f* since sin *θ* ≈ *θ* and cos *θ* ≈ 1 when *θ* is small (Fig. 2(c)). Moreover, the width of the off-zero peak, characterized by the difference of the frequencies at which the spectrum attains half of its peak value (see Fig. 2(c) for an illustration), is given by *D* = 2*πf/N*. In other words, the width is proportional to both the cell cycle frequency and cell cycle duration variability (in agreement with fig. S1(a)). In the case of bursty gene expression, the height of the off-zero peak is given by (see section S4 for the proof)

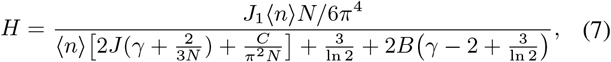

where *γ* = 2(1 − 4*pq*)*/*(1 + 2*pq*) is a function of *p*, and

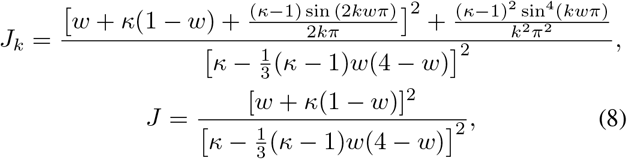

and 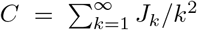 are all functions of *κ* and *w*. In the non-bursty case, the term involving *B* in Eq. (7) vanishes. Note that since *p* ≤ 1*/*2 is the probability that a molecule is allocated to one daughter and *q* = 1 − *p* is the probability of being allocated to the other daughter, it follows that 0 ≤ *γ* ≤ 2; hence the parameter *γ* is a dimensionless measure of the asymmetry of cell division. From Eq. (7), it is easy to see that *H* decreases with increasing *B* and decreasing *p* and *n*. This is in full agreement with the numerical results shown in Fig. 2(d),(e) which are computed using Eq. (2).

On the other hand, in bacteria and yeast, most mRNAs have very short half-lives, i.e. *η ≫* 1 (table S1). For such unstable gene products, we also derive a simplified expression of the power spectrum which is given in Methods. The width of the off-zero peak is still given by *D* = 2*πf/N* but the height is given by

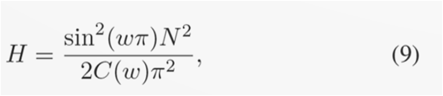

where 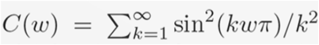 is a function of *w* (see section S5 for the proof). We emphasize that this formula is derived in the limit of *η* → ∞. In naturally occurring systems, the value of *η* for an unstable gene product usually does not exceed 100 (table S1) and hence applying this formula may lead to some errors, especially when *κ* is very close to 1. It is interesting to note that for unstable gene products, the height of the off-zero peak depends less on asymmetric division, gene expression mean, random bursting, and dosage compensation, whereas from Eq. (7), it is clear that the opposite is true for stable gene products. For both stable and unstable cases, as cell cycle duration variability becomes smaller (*N* increases), the off-zero peak becomes narrower and higher.

Recall that for quasi-symmetric cell division, sustained oscillations are the most regular when *w* is neither too small nor too large (Fig. 2(d)). For stable gene products with *η* ≪ 1, it follows from Eq. (7) that the maximal regularity is obtained when *w* ≈ 0.29 in the case of symmetric division, large gene expression mean, and no dosage compensation. For unstable gene products with *η ≫* 1, it follows from Eq. (9) that the maximal regularity is obtained when *w* = 0.5.

Note that our conclusions for the differences between the power spectra of (unstable) mRNA and (stable) protein are also confirmed by numerical simulations of a more complex model with both mRNA and protein descriptions (rather than an effective protein description with bursting dynamics as described in point 2 in the Model Specification section).

### Increasing the asymmetry of cell division induces a sharp change in the power spectrum for stable gene products

Our theory further reveals an important difference between symmetric (*p* = 1*/*2) and asymmetric (*p <* 1*/*2) cell division for stable gene products. When the gene expression mean is large, it follows from Eq. (7) that the height of the off-zero peak reduces to *H* ≈ *J*_1_*N/*6*π*^4^[2*J* (*γ* + 2*/*3*N*) + *C/π*^2^*N*]. For symmetric division, we have *γ* = 0 and thus the height depends on *N* quadratically as *H* ≈ *J*_1_*N* ^2^*/*2*π*^2^(4*π*^2^*J* + 3*C*) (fig. S2(a)). For asymmetric division, however, we have 0 *< γ* ≤ 2 and thus when cycle cycle duration variability is very small (*N ≫* 1), the height depends on *N* linearly as *H* ≈ *J*_1_*N/*12*π*^4^*Jγ* (fig. S2(b)). As cell division asymmetry becomes stronger, the height transitions sharply from the *N* ^2^ law to the *N* law (fig. S2(c)). This implies that for a certain cell cycle duration variability (fixed *N*), symmetric division leads to a much higher degree of regularity in oscillations for stable products than asymmetric division. In contrast, we find that for unstable gene products, there is no analogous transition because the height of the off-zero peak is always proportional to *N* ^2^ and is independent of *p* which controls the asymmetry of cell division (see Eq. (9)).

### Single-cell time traces can display higher-order harmonics of the cell cycle frequency

Interestingly, when cell cycle duration variability is very small, besides the peak at the cell cycle frequency *f*, the power spectrum also has peaks at integer multiples of *f* (Fig. 2(f) and fig. S2(a),(b)). This shows that besides the fundamental period of the mean doubling time *T* = 1*/f*, the system also has the hidden periods of *T/*2 and even *T/*3. Note that in Fig. 2(a),(c), we have not observed higher-order harmonics because the cell cycle duration variability is not sufficiently small (*N* is not sufficiently large). Similar peaks at higher-order harmonics have been previously reported for biochemical systems with feedback loops, due to the combination of intrinsic noise and nonlinearity in the law of mass action [41]. In the present model, the propensities of the reactions are all linear in molecule numbers and hence the hidden periods cannot be attributed to the same mechanism as in [41].

These hidden frequencies can be better understood using our analytical results. Recall that the spectral peak at *f* is controlled by the function *G*_1_(*ξ*) in the second term of Eq. (5). In fact, the remaining functions *G*_*k*_(*ξ*), *k* ≥ 2 in the second term control the peaks at higher-order harmonics *kf*. From Eq. (6), for stable gene products, the height and width of the spectral peak at the *k*th harmonic frequency are given by *J*_*k*_*H/J*_1_*k*^4^ and *k*^2^*D*, respectively, where *J*_*k*_ is given in Eq. (8). In particular, when *w* is very small or very large, we have *J*_*k*_ ≈ 1 for all *k* and hence the height of the spectral peak at *f* is 2^4^ = 16 times greater than that at 2*f* and 3^4^ = 81 times greater than that at 3*f*. Moreover, the width of the spectral peak at *f* is 2^2^ = 4 times lesser than that at 2*f* and 3^2^ = 9 times lesser than that at 3*f*. This is in full agreement with our simulation results which show that the peaks at higher-order harmonic frequencies are much lower and wider than the peak at the fundamental frequency (Fig. 2(f) and fig. S2(a),(b)).

For unstable gene products, similar phenomena are also observed but the characteristics of the higher-order spectral peaks are slightly different. Actually, the height and width of the spectral peak at the *k*th harmonic frequency are given by sin^2^(*kwπ*)*H/* sin^2^(*wπ*)*k*^4^ and *k*^2^*D*, respectively (see Methods for the proof). When *w* is very small or very large, we have sin^2^(*kwπ*)*/* sin^2^(*wπ*) ≈ *k*^2^ and hence the height of the spectral peak at *f* is 2^2^ = 4 times greater than that at 2*f* and 3^2^ = 9 times greater than that at 3*f*. Another interesting prediction is that for midway replication (*w* = 0.5), there are no peaks at even harmonics since the height at 2*kf* is zero (inset of Fig. 2(f)). As a result, we find that for midway replication, stable gene products yield higher peaks at higher-order harmonics, while for early or late replication, unstable gene products yield higher peaks at higher-order harmonics.

### Bifurcations between the four types of power spectra are observed as the gene product stability is varied

The dependence of oscillations on gene product stability *η* is much more complicated. Fig. 3(a)-(c) illustrate the height of the off-zero spectral peak as a function of *η, κ*, and *N* when the gene expression mean is large. Since type I spectra are monotonically decreasing, the height of the off-zero peak is set to be zero for convenience. For stable gene products with *η* ≪ 1, it is only possible to observe type I, II, and III spectra. It can be analytically shown that for the case of symmetric division, large gene expression mean, midway replication (*w* = 0.5), and no dosage compensation, stable gene products yield type I spectra when *N* ≤ 6, type II spectra when 7 ≤ *N* ≤ 28, and type III spectra when *N* ≥ 29 (see section S4 for the proof).

**Fig. 3.**
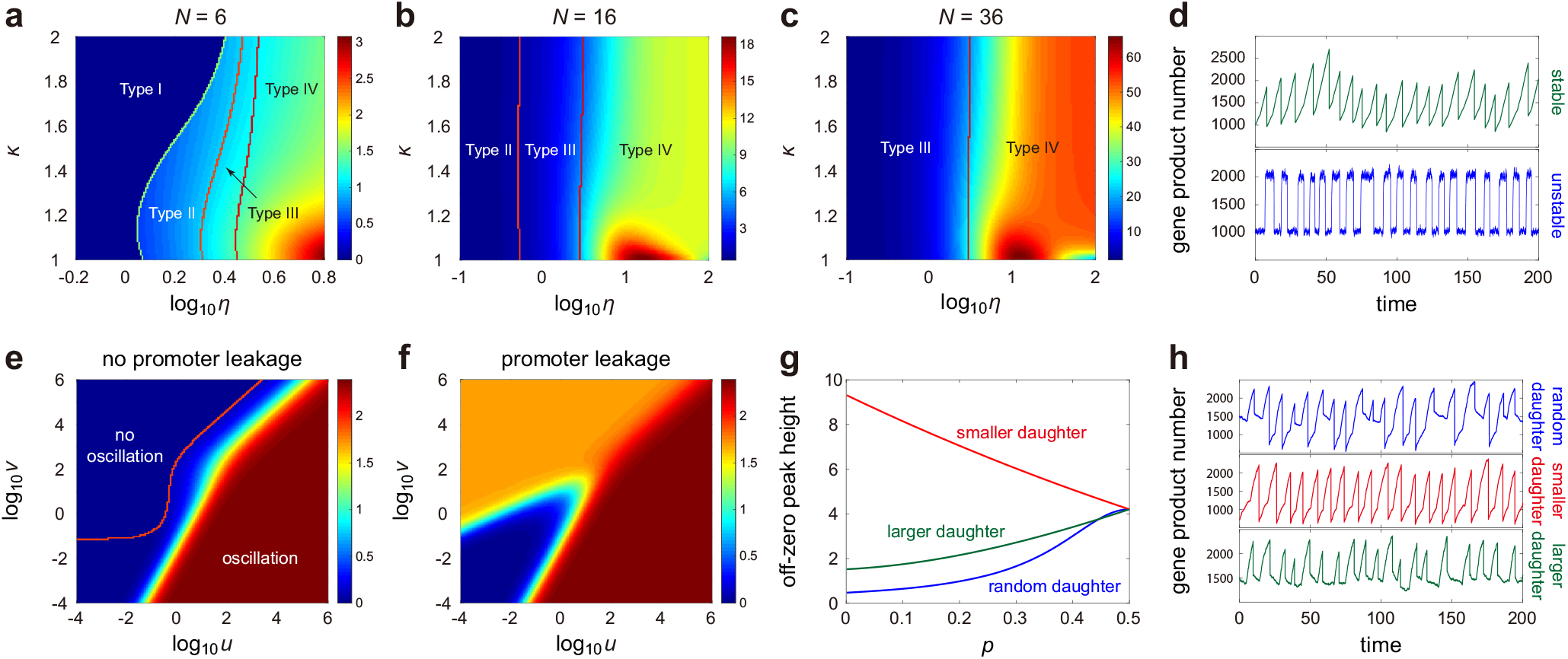
Further properties of the power spectrum. (**a**)-(**c**) Heat maps showing the dependence of the height of the off-zero peak (relative to the zero peak) on the parameters controlling gene product stability (*η*) and dosage compensation (*κ*). For stable gene products with *η* ≪ 1, it is only possible to observe type I-III spectra; type IV can be produced by unstable gene products with *η ≫* 1. Strong dosage compensation (*κ* = 1) in (b),(c) leads to maximal oscillation regularity at an intermediate gene product stability. (**d**) Typical trajectories for stable and unstable gene products. With the same gene expression mean, unstable products lead to more robust sustained oscillations than stable ones. (**e**) Heat map showing the dependence of the height of the off-zero peak (relative to the zero peak) on promoter switching rates *u* and *v* in the absence of promoter leakage. (**f**) Same as (e) but in the presence of promoter leakage. Fast promoter switching, leakage, and active gene state dominance enhance oscillation regularity. (**g**) Height of the off-zero peak (relative to the zero peak) versus the asymmetric allocation probability *p* for three types of tracking protocols at cell division: tracking one of the two daughters randomly (blue), tracking the smaller daughter (red), and tracking the larger daughter (green). **(h)** Typical trajectories for the three protocols generated using Gillespie’s algorithm. Note that (a)-(c) and (e)-(g) are obtained from numeric evaluation of Eq. (2). See section S11 for the technical details of this figure.

When *N* is small and *η* is not restricted to small values, all the four types of power spectra can be observed; as *η* increases, the system undergoes three stochastic bifurcations from type I to type II, then to type III, and finally to type IV (Fig. 3(a)); when *N* is moderate, type I spectra cannot occur; as *η* increases, the system undergoes two stochastic bifurcations from type II to type III, and then to type IV (Fig. 3(b) and fig. S1(b)); when *N* is large, both type I and II spectra fail to occur; as *η* increases, the system undergoes only one stochastic bifurcation from type III to type IV (Fig. 3(c)). These results show that sustained oscillations tend to occur when (i) *N* is small but *η* is large (unstable gene products with large cell cycle duration variability) or (ii) *N* is moderate or large. In the latter case of moderate or low cell cycle duration variability, oscillations can be observed for all values of *η* and hence are expected for both stable and unstable gene products.

The reason why unstable gene products yield more robust sustained oscillations can be understood as follows. For stable gene products, since active degradation is weak, molecule numbers increase approximately linearly with time and hence the time traces of gene product abundances appear like a sawtooth wave (upper panel of Fig. 3(d)). Due to cell cycle duration variability and noise due to partitioning at division, the expression levels at the end (or beginning) of each generation have large fluctuations. Such noise gives rise to the zero peak of the power spectrum, which explains why for small *η*, the spectra are of types I-III. In contrast, for unstable gene products, since the degradation rate is large, molecule numbers quickly reach a steady state and hence the time traces of gene product abundances appear like a square wave (lower panel of Fig. 3(d)). The two levels of the square wave correspond to the steady-state levels before and after replication. Once the expression level deviates from the steady-state value, the large degradation rate will help it relax to the steady state rapidly and hence the expression levels at the end (or beginning) of each generation have relatively small fluctuations. This explains why for unstable gene products with large *η*, the spectra are of type IV, i.e. do not have a peak at zero frequency, but rather the power is concentrated in a narrow bandwidth of frequencies close to the cell cycle frequency.

### Strong dosage compensation causes resonance-like behavior at intermediate gene product stability

The pattern of sustained oscillations is also influenced by dosage compensation. When dosage compensation is weak (*κ* is close to 2), the height of the off-zero spectral peak (relative to the zero peak) is an increasing function of *η* (Fig. 3(a)-(c)). In this case, the more unstable are gene products, the more capable they are of exhibiting regular stochastic oscillations. However, when dosage compensation is strong (*κ* is close to 1) and cell cycle duration variability is not too large, there is an optimal *η* such that the height is maximized (Fig. 3(b),(c)); oscillations are the most regular when *η* is around 15, which falls within the biological range of a typical mRNA in bacteria (table S1). The reason of this phenomenon can be understood as follows. Oscillations cannot be very regular for small *η* due to noise in cell cycle duration and partitioning at division, as discussed earlier. For large *η*, oscillations also cannot be very regular because when *κ* is close to 1, there is little change in the effective burst production rate across the cell cycle and hence the steady-state expression levels before and after replication (shown as the two levels of the square wave in Fig. 3(d)) will merge into one. As a result, when dosage compensation is strong, oscillations are the most regular at an intermediate *η* value.

### Fast promoter switching, leakage, and active gene state dominance enhance oscillation regularity

Finally, we investigate the effect of promoter switching on gene expression oscillations. Fig. 3(e),(f) illustrate the height of the off-zero spectral peak as a function of the promoter switching rates, *u* and *v*, in the absence and presence of promoter leakage, where promoter leakage means that there is a nonzero burst production rate when the gene is in the inactive state. For the case of no leakage, oscillations are manifest when the gene is mostly active, i.e. *u ≫ v*, and fail to occur when the gene is mostly inactive, i.e. *u* ≪ *v*, likely due to an exceptionally small gene expression mean. In the presence of leakage, however, we observe a strong oscillation when *u ≫ v* and a weaker oscillation when *u* ≪ *v*. Due to promoter leakage, even when the gene is mostly inactive, there is still a smaller but non-vanishing gene expression mean, which leads to the weaker oscillation observed. In addition, we find that the height of the off-zero peak is exceptionally small when promoter switching is very slow [42], i.e. *u, v* ≪ 1, regardless of whether there is promoter leakage or not. We stress here that while Fig. 3(e),(f) display the results for the inheritance mechanism, similar results also hold for the reset mechanism.

### The power spectrum in asymmetrically dividing cells is strongly influenced by the single-cell tracking protocol

In some previous papers [1, 43], to track a cell lineage, one of the two daughters was randomly selected at cell division with probability 1*/*2. However, for asymmetric cell division, the two daughters are of different sizes and another possible protocol is to track the smaller/larger daughter (such as the bud/mother cell in budding yeast) at cell division [3, 44]. Assuming well-mixing, the probability of a newborn cell receiving a gene product molecule is equal to the ratio of the volume of the newborn to the volume of the mother cell and hence on average the smaller daughter receives fewer gene products than the larger one. We remind the reader that in our model the probability for a molecule to be allocated to one daughter is *p* ≤ 1*/*2 and the probability of being allocated to the other is *q* = 1 − *p*. Hence it follows that the daughter with allocation probability *p <* 1*/*2 at division is the smaller daughter (Fig. 1(b)). Thus far we assumed that one of the two daughter cells is randomly tracked with probability 1*/*2 after cell division. Now we study two other tracking protocols, namely where we always follow the smaller or the larger daughter after cell division. For a cellular state (*N, i, j*) in effective cell cycle stage *N*, suppose that *i* and *j* record the gene states of the daughter copies that will be allocated to the smaller and larger daughter, respectively. If the smaller daughter is tracked after division, the cell will transition from a cellular state (*N, i, j*) in stage *N* to the cellular state (1, *i*) in stage 1 with rate *a*. If the larger daughter is tracked after division, the cell will transition from (*N, i, j*) to (1, *j*) with rate *a*. These considerations show that the transition diagram of cellular states should be modified as follows: if the smaller (larger) daughter is tracked at division, then the green (red) arrows in Fig. 1(d),(e) should be deleted and the transition rates of the red (green) arrows should be changed from *a/*2 to *a* (fig. S3).

In this case, the autocorrelation function has the same form as in Eq. (1), where *Q* = (*q*_*αβ*_) is the generator matrix of cellular state transitions (see fig. S3 for the transition diagram which has some colored arrows) and *W* = (*w*_*αβ*_) is another matrix obtained from *Q* by replacing the rates of orange and red (green) arrows by *pa* (*qa*) and subtracting *d* from the diagonal entries if the smaller (larger) daughter is tracked after cell division (see section S6 for the detailed expressions of each term).

To compare the three types of tracking protocols at cell division (tracking a random daughter, the smaller daughter, or the larger daughter), we illustrate the height of the off-zero spectral peak as a function of *p* for moderately unstable gene products with *η* = 2 (Fig. 3(g)). Clearly, the three tracking protocols give rise to the same oscillatory behavior for symmetric division. However, for asymmetric division, the smaller daughter tracking protocol yields a much higher off-zero peak than the other two protocols, implying the most regular oscillations. The reason for this observation is as follows. For unstable gene products, the molecule number relaxes to the steady-state value rapidly and hence the expression levels just before division are roughly the same for the three protocols. However just after division, the smaller daughter receives less molecules than the larger one. Hence it follows that smaller daughter tracking yields larger amplitudes of oscillations in the time traces than larger daughter tracking (Fig. 3(h)). Compared to random tracking, smaller/larger daughter tracking leads to less noisy oscillations, presumably since the latter introduces a deterministic element in the tracking process. This explains why random tracking leads generally to less robust oscillations than the other two tracking protocols (Fig. 3(g)). Note that while increasing cell division asymmetry (decreasing *p*) reduces the robustness of oscillations for random and larger daughter tracking, it leads to the opposite effect for small daughter tracking.

In fig. S4, we also investigate the differences between the power spectra obtained from the three tracking protocols as a function of the parameter *η* which is a measure of gene product stability. We find that the differences increase with gene product stability. This can be explained as follows. The abrupt change in the number of molecules =at division as we switch from a mother cell to a daughter cell is sensitive to the choice of protocol; these low frequency fluctuations contribute to the height of the zero peak which is sizeable for very stable products and fairly small for very unstable products (Fig. 3(a)-(c)). Hence the choice of protocol has most effect on the spectra of stable products.

### Experimental validation of the theory and its application to parameter inference

To test our theory, we apply it to study oscillations in single-cell gene expression data collected for *E. coli* in [2]. In this data set, the time course data of fluorescence intensity of a constitutively expressed yellow fluorescent protein was recorded every minute for 279 cell lineages over 70 generations using a mother machine under three different growth conditions (25°C, 27°C, and 37°C). At the three temperatures, there are a total of 65, 54, and 160 cell lineages measured, respectively. Based on such data, it is possible to estimate all the parameters involved in our model for each cell lineage. The medians of the estimated parameters for all cell lineages are listed in Table 2 and the distributions of the estimated parameters are given in figs. S5 and S6. In the following, we briefly describe the estimation procedures.

**Table 2.** The medians of the estimated parameters for all cell lineages at three different temperatures. The distributions of the estimated parameters can be found in figs. S5 and S6. The medians rather than the means are reported here since the estimation of the former is more robust than that of the latter, with respect to outliers.

|  | 25°C | 27°C | 37°C |
| --- | --- | --- | --- |
| $f$ (min<sup>-1</sup>) | 0.0148 | 0.0187 | 0.0307 |
| $T = 1/f$ (min) | 67.5676 | 53.4759 | 32.5733 |
| $N$ | 14 | 31 | 31 |
| $w$ | 0.2000 | 0.1450 | 0.1224 |
| $\kappa$ | 2.0621 | 2.0000 | 2.9412 |
| $\beta$ | 42.0719 | 15.4767 | 13.0344 |
| $\langle n \rangle$ | 99.2042 | 199.5930 | 287.8168 |
| $\rho_{\text{eff}}$ (min<sup>-1</sup>) | 0.1299 | 0.2303 | 0.6259 |
| $\kappa\rho_{\text{eff}}$ (min<sup>-1</sup>) | 0.2232 | 0.5319 | 1.3809 |
| $B$ | 4.7718 | 5.2099 | 4.8606 |

Since the protein is constitutively expressed [2], there is no promoter switching, i.e. *u* = *v* = 0. Moreover, since the protein is very stable, it is reasonable to assume that it has negligible degradation, i.e. *d* = *η* = 0 [2, 43]. Based on the time course data, we estimated the average power spectrum over all cell lineages at each temperature by means of the Wiener-Khinchin theorem (Fig. 4(a)), where we have normalized the spectrum so that *G*(0) = 1. Clearly, the average power spectra are of type II for all the three growth conditions. As the temperature increases, the position and height of the off-zero spectral peak both increase, implying more robust oscillations. The position of the off-zero peak is very close to the cell cycle frequency *f*, which can be easily estimated from the data of doubling times (Table 2).

**Fig. 4.**
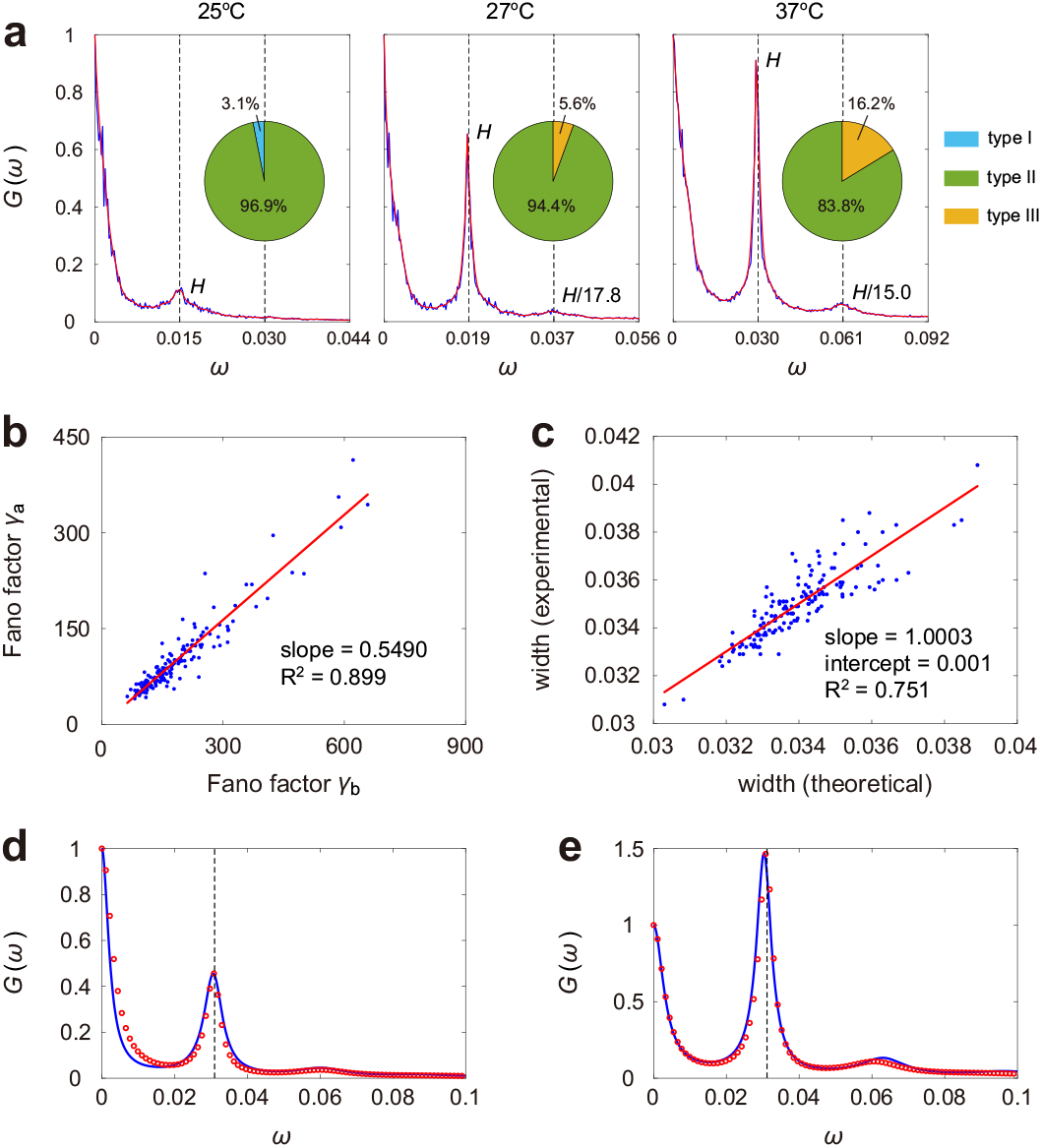
Analysis of single-cell time course data of fluorescence intensities in *E. coli* published in [2]. **(a)** Average power spectra over all cell lineages at three different temperatures (blue curves) and their smoothed approximations (red curves) using the Gaussian filter. The average spectra are estimated by means of the Wiener-Khinchin theorem, where 65, 54, and 160 cell lineages are averaged over, respectively, for the three growth conditions. The power spectrum for each cell lineage is also estimated by fitting the time course data with an AR model. The pie charts display the percentages of various types of power spectra for all cell lineages. **(b)** Comparison between the Fano factor *γ*_*b*_ of the fluorescence intensities just before division and the Fano factor *γ*_*a*_ of the fluorescence intensities just after division for all cell lineages at 37°C. **(c)** Comparison between the widths of the experimental power spectra obtained using the AR model technique and the theoretical power spectra determined using the estimated parameters for all cell lineages at 37°C. **(d)**,**(e)** Comparison between the experimental (blue curve) and theoretical (red circles) power spectra for two typical cell lineages at 37°C.

We find that the doubling time data for all cell lineages are well fitted by an Erlang distribution; the parameter *N* can then be estimated as the inverse of the coefficient of variation squared of this distribution. The medians of the estimated *N* for the three growth conditions are 14, 31, and 31, respectively. It can be seen that cells cultured at 27°C and 37°C have much smaller cell cycle duration variability than those cultured at 25°C. Our theory demonstrates that smaller cell cycle duration variability gives rise to a higher off-zero spectral peak (Fig. 2(c)). This explains why the off-zero peak is significantly higher at higher temperatures (Fig. 4(a)). Our theoretical result also predicts that when *N* is large, the power spectrum has peaks at higher-order harmonics of the cell cycle frequency (Fig. 2(f)). This is in excellent agreement with our data analysis, which shows that the average power spectra at 27°C and 37°C have an apparent peak at the second harmonic frequency. Interestingly, the ratio of the heights of the spectral peaks at *f* and 2*f* is estimated to be 17.8 and 15.0 for the two temperatures, both of which are very close to the theoretical value of 2^4^ = 16 predicted by our theory for stable gene products when *w* is small.

To perform a more detailed analysis, we also estimated the power spectrum for each cell lineage by fitting the time course data with an autoregressive (AR) model, which is a standard model in time series analysis [45], with the order of the AR model being determined by minimizing the Akaike information criterion (see section S7 for details). Our theoretical results show that for stable gene products, only type I, II, and III spectra can be observed (Fig. 3(a)-(c)). This is in full agreement with our data analysis with the percentages of the three types of power spectra being illustrated by the pie charts in Fig. 4(a). Clearly, type II spectra are dominant for all the three growth conditions. For cells at 25°C, only types I and II are observed; for cells at higher temperatures, only types II and III are observed. The percentage of type III spectra is higher for cells at 37°C than cells at 27°C.

Since the burst production rate increases from *ρ*_eff_ to *κρ*_eff_ upon replication, we can fit the data (recorded per minute) between two cell division times by the following mean-field approximation:

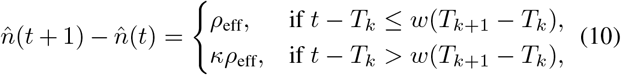

where *T*_*k*_ is the *k*th cell division time and *t* is an arbitrary time point (in minute) between two consecutive division times *T*_*k*_ and *T*_*k*+1_. Here we use a piecewise linear function to approximate the time series of expression levels between two division events. The first (second) part of the piecewise linear function approximates the time series before (after) replication and thus should have an average slope of *ρ*_eff_ (*κρ*_eff_). By fitting the time course data *n*(*t*) with the approximation 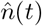 given in Eq. (10), we obtained the least squares estimates of *w* and *κ* for each cell lineage by minimizing the distance 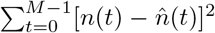 between the two, where *M* is the number of time points for each cell lineage (see section S8 for details). The medians of the estimated *w* for the three growth conditions are 0.20, 0.15, and 0.12, respectively, which decrease with temperature. This is possibly because at higher temperatures, the growth rate of *E. coli* cells is faster and thus replication would occur more continuously, leading to a shift in *w* towards zero [46]. In addition, the medians of the estimated *κ* are close to 2 for all the three growth conditions (Table 2), implying weak dosage compensation.

The remaining parameters to be estimated are *ρ*_eff_ and *B*, where *B* is the mean translational burst size, i.e. the average number of protein molecules produced per mRNA lifetime. To estimate them, recall that *ρ*_eff_*B* represents the mean number of protein molecules produced per unit time. However, what was measured in the data set is the fluorescence intensity of protein molecules, instead of the real copy number. Hence, it is crucial to determine the proportionality constant between fluorescence intensities and copy numbers. To do this, we note that under the assumption of symmetric binomial partitioning at division, the expression levels just before and just after a particular cell division time are coupled by (see section S9 for the proof)

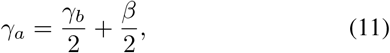

where *γ*_*b*_ (*γ*_*a*_) is the Fano factor (the variance divided by the mean) of the fluorescence intensities just before (after) division and *β* is the fluorescence intensity per protein molecule. Note that both *γ*_*b*_ and *γ*_*a*_ can be estimated for each cell lineage. To test this relationship, we show *γ*_*a*_ as a function of *γ*_*b*_ for all cell lineages at the three temperatures (see Fig. 4(b) for cells at 37°C and fig. S7(a) for cells at lower temperatures), from which we observed a strong linear relationship with a high *R*^2^ and a slope close to 0.5. Then the proportionality constant *β* for a given temperature can be estimated as *β* = 2 ⟨*γ*_*a*_⟩ − ⟨*γ*_*b*_⟩, where the angled brackets denote the sample means over all cell lineages at that temperature. The estimated *β* for the three growth conditions are 42.1, 15.5, and 13.0, respectively. In particular, our analysis shows that the fluorescence intensity per protein molecule decreases with temperature, which is likely because lower temperatures are more conducive to the correct folding of the fluorescent protein [47].

From the fluorescence intensity data and the inferred *β*, it is easy to estimate the gene expression mean ⟨*n*⟩ for each cell lineage (Table 2). Note that cells at 27°C and 37°C have similar cell cycle duration variability with *N* = 31 but the latter yield a higher off-zero spectral peak. This is because cells at 37°C have a larger gene expression mean, which augments the height of the off-zero spectral peak. Moreover, since *ρ*_eff_, *B, w, κ*, and ⟨*n*⟩ are related by Eq. (3) and we have estimated *w* and *κ*, we can obtain the estimate of *ρ*_eff_*B*. Finally, we estimated *ρ*_eff_ and *B* separately by equating the heights of the off-zero peak of the experimental power spectrum obtained using the AR model technique and the theoretical power spectrum determined by Eq. (2) using all the estimated parameters. From Table 2, we can see that the mean burst frequency (average number of bursts produced per unit time) *ρ*_eff_ increases with temperature approximately linearly but the mean burst size *B* is not temperature dependent.

Thus far we have shown how, by means of the theoretical expressions of the power spectrum and the gene expression mean, we can estimate all the model parameters for each cell lineage from the time course data. The accuracy of these estimates is verified in two different ways. First, in Fig. 4(d),(e), we show a good agreement between the experimental power spectrum and the theoretical power spectrum evaluated using the inferred model parameters for two typical cell lineages. As a second test of the accuracy of parameter inference, we compare the widths of the experimental and theoretical power spectra for all cell lineages (see Fig. 4(c) for cells at 37°C and fig. S7(b) for cells at lower temperatures). It can be seen from Fig. 4(c) that the widths for the two spectra show a strong linear relationship with a slope close to 1, a negligible intercept, and an *R*^2^ of 0.75. We stress that while the model parameters were not estimated from the full spectrum curve but only from the height of the off-zero peak, the full theoretical spectrum matches the experimental spectrum reasonably well.

## Discussion

In this work, we have investigated the frequency decomposition of the copy number fluctuations of a gene product (mRNA or protein) within a cell lineage, by deriving expressions for the power spectrum of fluctuations in a detailed stochastic model of gene expression. This model takes into account the salient experimental observations of intracellular dynamics including promoter switching, transcriptional and translational bursting, variability in the duration of the cell cycle, variability in the gene copy number due to replication, the copying or resetting of the gene state during replication, gene dosage compensation, and partitioning of molecules due to symmetric or asymmetric cell division.

Our study differs from previous ones in four main respects: (i) our model is more grounded in biological reality than other models in the literature due to the large number of subcellular and cellular processes that it accounts for, as mentioned above; (ii) a number of studies have derived the distribution of molecule numbers or more commonly the moments for models that have some similarity to ours [15–19, 48]; however in contrast, here we derive expressions for the power spectrum which provide information about the frequency content of lineage data. This type of analysis has previously been only reported for models of non-growing cells [20, 49]; (iii) rather than a parameter inference based on the matching of the moments or the distribution of molecule numbers calculated from the data to those of a stochastic model [50], we showcase a power spectrum based parameter inference method; (iv) our model takes into account the details of the experimental protocol used for tracking cells across a lineage.

Our novel theory provides expressions for the height and width of the off-zero spectral peak (with its position close to the cell cycle frequency) as a function of all rate parameters in the model. The ratio of the height of this peak to the power at zero frequency provides a means to classify the power spectra into two main types: (i) the spectra with the ratio less than 1 (types I and II) and (ii) the spectra with the ratio greater than 1 (types III and IV). The periodicity in molecule number variation induced by cell division dominates over subcellular noise for (ii) while the reverse is the case for (i). Type I is further differentiated from type II by specifying that in the former, there is only one peak at zero frequency whereas in the latter there is a dominant peak at zero frequency and a lesser one at approximately the cell cycle frequency. Similarly type III is further differentiated from type IV by specifying that in the former, there is a dominant peak at approximately the cell cycle frequency and a lesser one at zero frequency while in the latter there is only one peak at approximately the cell cycle frequency. The theory also predicts that while the spectra for fast decaying (unstable) gene products can be of all four types, the spectra for slow decaying (stable) gene products can only be of types I-III.

Our analysis of experimental data for *E. coli* shows that the type of spectra of single-cell trajectories depends on the temperature: lower temperatures favour type I and II spectra while higher temperatures favour type II and III spectra. Overall the most common spectrum was Type II implying that for many cells, the “forces” inducing periodicity of molecule numbers are typically slightly less strong than the “forces” inducing subcellular noise; the strength of the latter increases with decreasing temperature. None of the 279 cell lineages had a type IV spectrum for protein fluctuations, in accordance with the theoretical result that stable gene products cannot display such a spectrum.

Our theory made a number of other testable predictions: (i) the height of the off-zero spectral peak (relative to that of the zero peak) increases with decreasing cell cycle duration variability and increasing mean expression levels, while the width of the off-zero peak is proportional to the cell cycle frequency and to the cell cycle duration variability; (ii) if the cell cycle duration variability is small enough then the spectra display peaks at higher-order harmonics of the cell cycle frequency. For stable proteins, the height of the spectral peak at the second harmonic (twice the cell cycle frequency) is 16 times larger than that at the cell cycle frequency for early or late replication. Both predictions were confirmed by analysis of lineage data for *E. coli*.

Furthermore our theory made numerous other predictions which require the design of new experiments and which cannot be tested using current data. A brief summary of these predictions is as follows: (i) the dependence of the height of the off-zero spectral peak on parameters is very different for gene products that are stable, e.g. most proteins, compared to gene products that are unstable, e.g. most mRNAs. For stable gene products, the height of the off-zero peak decreases with increasing mean burst size, decreasing mean molecule number, and increasing asymmetry of cell division. Furthermore the height is maximal for replication occurring (almost) a third of the way through the cycle. In contrast for unstable gene products, the height of the off-zero peak is maximal for replication occurring midway through the cycle but is (almost) independent of asymmetric division, gene expression mean, random bursting, and dosage compensation. Independent of stability, fast promoter switching enhances the height of the off-zero peak (relative to that of the zero peak) whereas slow switching has the opposite effect; (ii) the strength of gene dosage compensation is reflected in how the spectrum varies with gene product stability. For weak dosage compensation, molecules which have a larger decay rate also exhibit a spectrum with a higher off-zero peak. For strong dosage compensation, molecules which have a decay rate that is neither too large nor too small are the ones which exhibit the most pronounced off-zero peaks; (iii) the power spectrum in asymmetrically dividing cells is strongly influenced by the choice of single-cell tracking protocol. In particular the spectra obtained from following the smaller daughter after division have a significantly higher off-zero peak than those obtained from following the larger daughter or from random tracking. This last prediction is likely the easiest of the three to check by redoing the cell tracking analysis in budding yeast which displays asymmetric cell division. Predictions (i) and (ii) ideally require the ability to obtain live-cell fluorescence data for mRNA over tens of generations and for different types of mRNA with widely varying decay rates, which are presently not easy to obtain. The development of such methods, particularly those with minimal perturbation of transcription and translation, is still in an active area of research [4].

We have also showcased the use of the power spectrum to determine the values of all the rate parameters in our model from cell lineage data. Two strengths of our inference procedure are (i) we do not need an experimental determination of the relationship between total fluorescence intensity and molecule number; this is determined automatically from the relationship between fluctuations in the fluorescence intensity just before and just after division; (ii) our analysis takes into account noise due to partitioning and variability in the cell cycle duration which has recently been shown to be crucial to obtain an accurate estimation of the burst frequency and the burst size [19]. Our analysis confirms there is no or very weak dosage compensation in *E. coli* in agreement with previous studies [46], i.e. at replication, the expression increases roughly two-fold, in agreement with an expected doubling of the number of gene copies. Furthermore we find that for constitutive expression, while the translational burst frequency increases approximately linearly with temperature, the translational burst size is temperature invariant. Previous studies were conducted at one temperature [5] and hence could not quantify the thermal dependence of gene expression parameters.

Concluding we have performed an exact frequency analysis of mRNA and protein fluctuations in a detailed model of gene circuit elements central to gene expression control. As we have shown, there is a wealth of information about subcellular processes hidden in the frequency content of mRNA and protein fluctuations within a cell lineage, and we hope our results will further stimulate work in this area.

## Methods

### Oscillations for diploid cells

Suppose that we have computed the analytical solutions of the autocorrelation function *R*(*t*) and the power spectrum *G*(*ξ*) for haploid cells. For diploid cells, we assume that the two alleles act independently of each other. Under the assumption of allelic expression independence, let *n*(*t*) and *n*′(*t*) denote the expression levels of the two alleles at time *t*, which are independent and identically distributed. Then for diploid cells, the autocorrelation function is given by

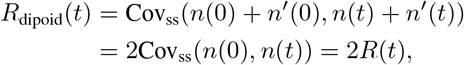

and thus the power spectrum is given by

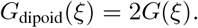

Since the two functions only different by a constant for haploid and diploid cells, they lead to the same oscillatory behavior. Thus we only need to focus on haploid cells in what follows.

### Master equation describing cell-cycle dependent stochastic gene expression

Recall that before gene replication, each cellular state can be represented as *α* = (*r, i*), where *r* is the cell cycle stage and *i* is the gene state of the mother copy; after gene replication, each cellular state can be represented as *α* = (*r, i, j*), where *i* and *j* are the gene states of the two daughter copies. Let *π*_*α*_ denote the probability of observing cellular state *α*. The evolution of the cellular state dynamics is then governed by the following master equation (see section S1 for the master equation written using the original model parameters):

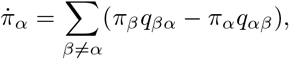

where *q*_*αβ*_ is the transition rate from cellular state *α* to cellular state *β*. The transition diagrams for all cellular states under the inheritance and reset mechanisms are illustrated in Fig. 1(d),(e), respectively. The microstate of the gene of interest can be represented by the ordered pair (*α, n*), where *α* is the cellular state and *n* is the copy number of the gene product. At division, the cell will transition from some microstate (*N, i, j, n*) in cell cycle stage *N* to another microstate (1, *i, m*) or (1, *j, m*) in cell cycle stage 1 with *m* ≤ *n*. To model asymmetric cell division, we need to distinguish the following two types of transitions between cellular states:

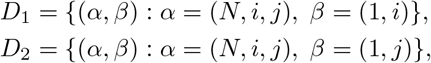

where *α* is a cellular state in cell cycle stage *N* and *β* is a cellular state in cell cycle stage 1. For any pair of cellular states (*α, β*) in *D*_1_ or *D*_2_, the transition rate from *α* to *β* is given as follows:

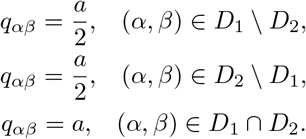

The cellular state transitions in *D*_1_ \ *D*_2_, *D*_2_ \ *D*_1_, and *D*_1_ ∩ *D*_2_ are marked by the red, green, and orange arrows in Fig. 1(d),(e), respectively. Due to asymmetric binomial partitioning at cell division, the transition rate from microstate (*α, n*) to (*β, m*) is given as follows:

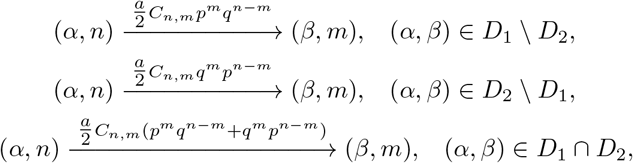

where *C*_*n,m*_ = *n*!*/m*!(*n* − *m*)! is the combinatorial number and *p* is the probability of asymmetric binomial partitioning. Let *p*_*α,n*_ denote the probability of being in microstate (*α, n*). Then the evolution of stochastic gene expression dynamics is governed by the following master equation:

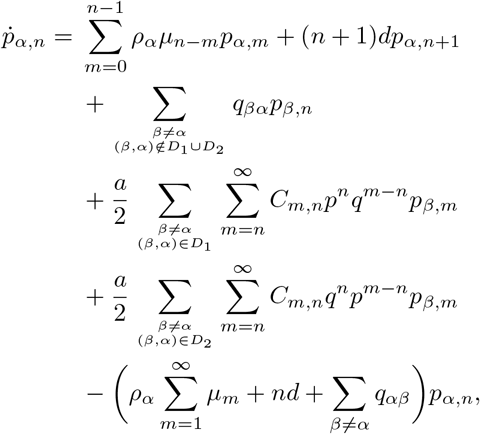

where *ρ*_*α*_ is the total burst production rate (for the mother copy or for the two daughter copies) in cellular state *α, µ*_*n*_ is the burst size distribution of the gene product, *d* is the degradation rate of the gene product, and *q*_*αβ*_ is the transition rate from cellular state *α* to cellular state *β*. Since there are only two gene states, there are only five possible values for *ρ*_*α*_, which are given by

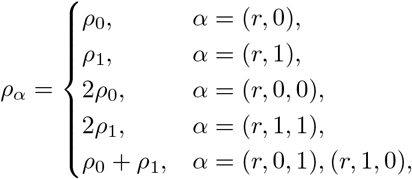

where *ρ*_0_ and *ρ*_1_ are the burst product rate (for a single gene copy) in the inactive and active gene states, respectively.

### Analytical solutions for the autocorrelation function and the power spectrum

Here we only consider the bursty case. The results in the non-bursty case can be derived in the same way (see section S2 for details). Let *α*(*t*) denote the state of the gene and let *n*(*t*) denote the copy number of the gene product in an individual cell at time *t*, respectively. To proceed, let

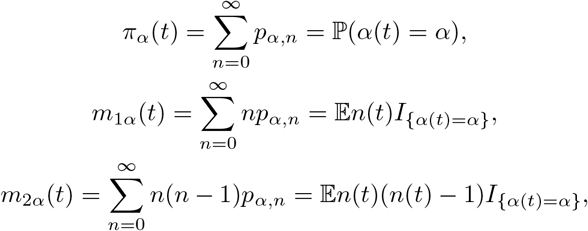

be the first three unnormalized factorial moments of gene product abundances in cellular state *α*, where *I*_*A*_ is the indicator function of the set *A*. Then the row vectors *π* = (*π*_*α*_), *m*_1_ = (*m*_1*α*_), and *m*_2_ = (*m*_2*α*_) satisfy the following differential equations (see section S2 for the proof):

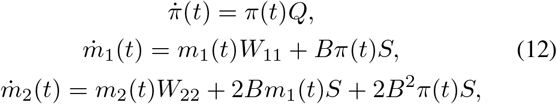

where *S* = diag(*ρ*_*α*_) is the diagonal matrix whose diagonal entries are the burst production rates in all cellular states and *W*_11_ and *W*_22_ are two matrices defined as

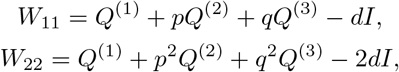

where 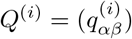 are three matrices defined as

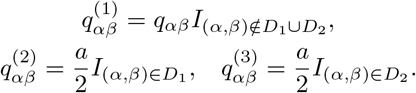

From Eq. (12), at the steady state, the first and second factorial moments are given by

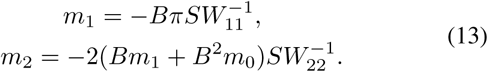

Therefore, the steady-state gene expression mean is given by

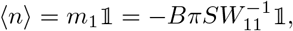

where 𝟙 denotes the column vector whose components are all 1.

Since Eq. (12) is a set of linear differential equations, its time-dependent solution is given by

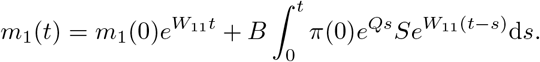

From now on, we assume that the system has reached the steady state. Given the initial cellular state *α*(0) = *α* and initial copy number *n*(0) = *n* of the gene product, it follows that

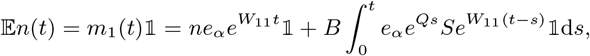

where *e*_*α*_ denotes the row vector whose *α*th component is 1 and all other components are 0. This clearly shows that

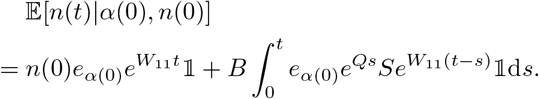

Therefore, at the steady state, we have (see section S2 for the proof)

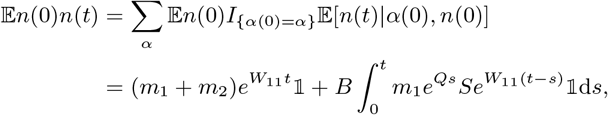

where *m*_1_ and *m*_2_ are the steady-state first and second factorial moments given in Eq. (13). Since the autocorrelation function is defined as *R*(*t*) = 𝔼*n*(0)*n*(*t*) − 𝔼*n*(0) 𝔼*n*(*t*), we finally obtain the explicit expression of the autocorrelation function, which is given in Eq. (1).

We next express *R*(*t*) more explicitly in terms of simple functions. To this end, we assume that all eigenvalues of *Q*, as well as all eigenvalues of *W*_11_, are mutually distinct (in fact, any matrix can be approximated by such matrices to any degree of accuracy). Let *λ*_0_, *λ*_1_, …, *λ*_*K−*1_ be all eigenvalues of *Q* and let *λ*_*K*_, *λ*_*K*+1_, *…, γ*_2*K−*1_ be all eigenvalues of *W*_11_. By the Perron-Frobenius theorem, one eigenvalue of the generator matrix *Q* must be zero and other eigenvalues must have negative real parts. Similarly, all eigenvalues of *W*_11_ must have negative real parts, which implies that

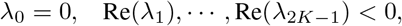

where Re(*x*) denotes the real part of *x*. From Eq. (1), it can be deduced that the correlation function is a linear combination of exponential functions:

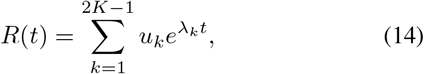

where *u*_*k*_ are suitable constants. Taking the derivatives on both sides of Eqs. (1) and (14) and evaluating at *t* = 0 yield

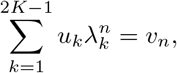

where

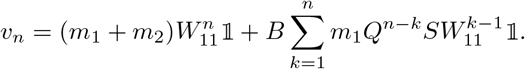

This equation can be rewritten in matrix form as

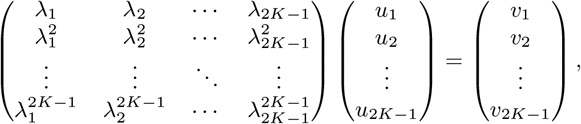

where the matrix *V* on the left-hand side is a Vandermonde matrix. This shows that

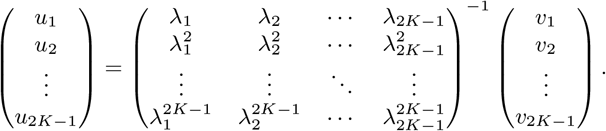

To proceed, recall that the inverse of the Vandermonde matrix is given by *V* ^*−*1^ = (*b*_*kl*_), where

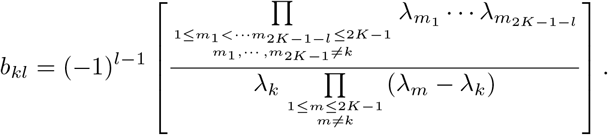

Therefore, the coefficients *u*_*k*_ are given by

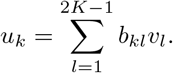

Finally, the autocorrelation function can be written as

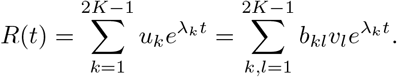

Taking the Fourier transform of the autocorrelation function gives the power spectrum, which is given in Eq. (2). The simplification of the gene expression mean and the power spectrum in the regime of fast promoter switching can be found in sections S2-S5.

### Power spectrum for stable gene products

For stable gene products with *η* ≪ 1, the power spectrum can be simplified as in Eq. (5), where

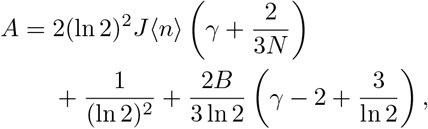

if gene expression is bursty,

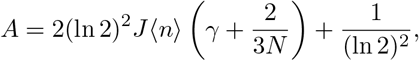

if gene expression is non-bursty, and

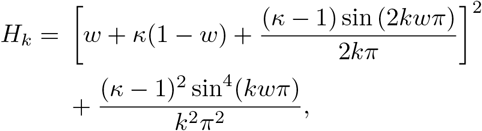

is a function of *κ* and *w*, where *J* is given in Eq. (8).

### Power spectrum for unstable gene products

For unstable gene products with *η* ≪ 1, when cell cycle duration variability is not too large, the power spectrum can be written more explicitly as (see section S5 for the proof)

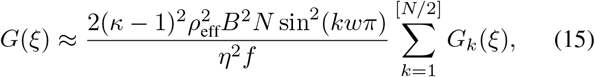

where

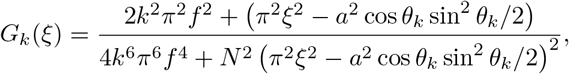

with *θ*_*k*_ = 2*kπ/N*. Here the function *G*_1_(*ξ*) controls the first off-zero spectral peak and the functions *G*_*k*_(*ξ*), *k* ≥ 2 control the peaks at higher-order harmonic frequencies. In analogy to the case of stable gene products, the position of the off-zero peak is given by *ξ* = (*a/π*) cos *θ*_1_ sin(*θ*_1_*/*2) *< aθ*_1_*/*2*π* = *a/N*, which is smaller than the cell cycle frequency *f*. When *N ≫* 1, the peak position is approximately equal to *f* since sin *θ* ≈ *θ* and cos *θ* ≈ 1 when *θ* is small. Moreover, the width of the off-zero peak is given by *D* = 2*πf/N* and the height of the off-zero peak is given in Eq. (9). From Eq. (15), the height and width of the spectral peak at the *k*th harmonic frequency are given by sin^2^(*kwπ*)*/* sin^2^(*wπ*)*k*^4^*H* and *k*^2^*D*, respectively. In particular, when cell cycle duration variability is small, the height of the spectral peak at *f* is 2^4^ sin^2^(*wπ*)*/* sin^2^(2*wπ*) times greater than that at 2*f* and 3^4^ sin^2^(*wπ*)*/* sin^2^(3*wπ*) times greater than that at 3*f*. Moreover, the width of the spectral peak at *f* is 2^2^ = 4 times lesser than that at 2*f* and 3^2^ = 9 times lesser than that at 3*f*.

## Acknowledgements

We thank Ruben Perez-Carrasco and Irina Kalita for stimulating discussions.

## Author contributions

R. G. conceived the original idea. C. J. performed the theoretical derivations and numerical simulations. C. J. and R. G. interpreted the theoretical results, analyzed the experimental data, and jointly wrote the manuscript.

## Data and materials availability

All data needed to evaluate the conclusions in the paper are present in the paper and in [2].

