## Supplementary information for "Frequency domain analysis of fluctuations of mRNA and protein copy numbers within a cell lineage: theory and experimental validation"

### Section S1. Master equation describing cell-cycle dependent stochastic gene expression

Recall that before gene replication, each cellular state can be represented as $\alpha = (r, i)$, where $r$ is the cell cycle stage and $i$ is the gene state of the mother copy; after gene replication, each cellular state can be represented as $\alpha = (r, i, j)$, where $i$ and $j$ are the gene states of the two daughter copies. Let $\pi_\alpha$ denote the probability of observing cellular state $\alpha$. The evolution of the cellular state dynamics is then governed by the following master equations. For $r = 1$, we have

$$\begin{aligned}
\dot{\pi}_{1,0} &= a(\pi_{N,0,0} - \pi_{1,0}) + \frac{a}{2}(\pi_{N,0,1} - \pi_{1,0}) + \frac{a}{2}(\pi_{N,1,0} - \pi_{1,0}) + (v\pi_{1,1} - u\pi_{1,0}), \\
\dot{\pi}_{1,1} &= a(\pi_{N,1,1} - \pi_{1,1}) + \frac{a}{2}(\pi_{N,0,1} - \pi_{1,1}) + \frac{a}{2}(\pi_{N,1,0} - \pi_{1,1}) + (u\pi_{1,0} - v\pi_{1,1}),
\end{aligned} \tag{1}$$

where $a$ is the transition rate from a cell cycle stage to the next, which is assumed to be the same for all cell cycle stages, the first three terms on the right-hand side correspond to cell division, and the fourth term corresponds to gene switching of the mother copy. For $2 \leq r \leq N_0$, we have

$$\begin{aligned}
\dot{\pi}_{r,0} &= a(\pi_{r-1,0} - \pi_{r,0}) + (v\pi_{r,1} - u\pi_{r,0}), \\
\dot{\pi}_{r,1} &= a(\pi_{r-1,1} - \pi_{r,1}) + (u\pi_{r,0} - v\pi_{r,1}),
\end{aligned}$$

where the first term corresponds to cell cycle progression and the second term corresponds to gene switching of the mother copy. For $r = N_0 + 1$, under the inheritance mechanism, we have

$$\begin{aligned}
\dot{\pi}_{N_0+1,0,0} = {} & a(\pi_{N_0,0} - \pi_{N_0+1,0,0}) + (v\pi_{N_0+1,0,1} - gu\pi_{N_0+1,0,0}) \\
& + (v\pi_{N_0+1,1,0} - gu\pi_{N_0+1,0,0}), \\
\dot{\pi}_{N_0+1,1,1} = {} & a(\pi_{N_0,1} - \pi_{N_0+1,1,1}) + (gu\pi_{N_0+1,0,1} - v\pi_{N_0+1,1,1}) \\
& + (gu\pi_{N_0+1,1,0} - v\pi_{N_0+1,1,1}), \\
\dot{\pi}_{N_0+1,0,1} = {} & -a\pi_{N_0+1,0,1} + (gu\pi_{N_0+1,0,0} - v\pi_{N_0+1,0,1}) + (v\pi_{N_0+1,1,1} - gu\pi_{N_0+1,0,1}), \\
\dot{\pi}_{N_0+1,1,0} = {} & -a\pi_{N_0+1,1,0} + (gu\pi_{N_0+1,0,0} - v\pi_{N_0+1,1,0}) + (v\pi_{N_0+1,1,1} - gu\pi_{N_0+1,1,0});
\end{aligned}$$

under the reset mechanism we have

$$\begin{aligned}
\dot{\pi}_{N_0+1,0,0} &= a(\pi_{N_0,0}+\pi_{N_0,1}-\pi_{N_0+1,0,0})+(v\pi_{N_0+1,0,1}-gu\pi_{N_0+1,0,0})\\
&\quad+(v\pi_{N_0+1,1,0}-gu\pi_{N_0+1,0,0}),\\
\dot{\pi}_{N_0+1,1,1} &= -a\pi_{N_0+1,1,1}+(gu\pi_{N_0+1,0,1}-v\pi_{N_0+1,1,1})+(gu\pi_{N_0+1,1,0}-v\pi_{N_0+1,1,1}),\\
\dot{\pi}_{N_0+1,0,1} &= -a\pi_{N_0+1,0,1}+(gu\pi_{N_0+1,0,0}-v\pi_{N_0+1,0,1})+(v\pi_{N_0+1,1,1}-gu\pi_{N_0+1,0,1}),\\
\dot{\pi}_{N_0+1,1,0} &= -a\pi_{N_0+1,1,0}+(gu\pi_{N_0+1,0,0}-v\pi_{N_0+1,1,0})+(v\pi_{N_0+1,1,1}-gu\pi_{N_0+1,1,0}),
\end{aligned}$$

where the first term corresponds to gene replication and the second and third terms correspond to gene switching of the two daughter copies. Finally, for $N_0+2\leq r\leq N$, we have

$$\begin{aligned}
\dot{\pi}_{r,0,0} &= a(\pi_{r-1,0,0}-\pi_{r,0,0})+(v\pi_{r,0,1}-gu\pi_{r,0,0})+(v\pi_{r,1,0}-gu\pi_{r,0,0}),\\
\dot{\pi}_{r,1,1} &= a(\pi_{r-1,1,1}-\pi_{r,1,1})+(gu\pi_{r,0,1}-v\pi_{r,1,1})+(gu\pi_{r,1,0}-v\pi_{r,1,1}),\\
\dot{\pi}_{r,0,1} &= a(\pi_{r-1,0,1}-\pi_{r,0,1})+(gu\pi_{r,0,0}-v\pi_{r,0,1})+(v\pi_{r,1,1}-gu\pi_{r,0,1}),\\
\dot{\pi}_{r,1,0} &= a(\pi_{r-1,1,0}-\pi_{r,1,0})+(gu\pi_{r,0,0}-v\pi_{r,1,0})+(v\pi_{r,1,1}-gu\pi_{r,1,0}),
\end{aligned}$$

where the first term corresponds to cell cycle progression and the second and third terms correspond to gene switching of the two daughter copies. The transition diagrams for all cellular states under the inheritance and reset mechanisms are illustrated in Fig. 1(d),(e) in the main text, respectively.

The microstate of the gene of interest can be represented by the order pair $(\alpha,n)$, where $\alpha$ is the cellular state and $n$ is the copy number of the gene product. At division, the cell will transition from some microstate $(N,i,j,n)$ in cell cycle stage $N$ to another microstate $(1,i,m)$ or $(1,j,m)$ in cell cycle stage 1 with $m\leq n$. To model asymmetric cell division, we need to distinguish the following two types of transitions between cellular states:

$$\begin{aligned}
D_1 &= \{(\alpha,\beta):\alpha=(N,i,j),\ \beta=(1,i)\},\\
D_2 &= \{(\alpha,\beta):\alpha=(N,i,j),\ \beta=(1,j)\},
\end{aligned}$$

where $\alpha$ is a cellular state in cell cycle stage $N$ and $\beta$ is a cellular state in cell cycle stage 1. For any pair of cellular states $(\alpha,\beta)$ in $D_1$ or $D_2$, the transition rate from $\alpha$ to $\beta$ is given as follows:

$$\begin{aligned}
q_{\alpha\beta} &= \frac{a}{2},\quad (\alpha,\beta)\in D_1\setminus D_2,\\
q_{\alpha\beta} &= \frac{a}{2},\quad (\alpha,\beta)\in D_2\setminus D_1,\\
q_{\alpha\beta} &= a,\quad (\alpha,\beta)\in D_1\cap D_2.
\end{aligned}$$

The cellular state transitions in $D_1\setminus D_2$, $D_2\setminus D_1$, and $D_1\cap D_2$ are marked by the red, green, and orange arrows in Fig. 1(d),(e) in the main text, respectively. Due to asymmetric binomial partitioning at cell division, the transition rate from microstate $(\alpha,n)$ to $(\beta,m)$ is given as follows:

$$\begin{aligned}
(\alpha,n) &\xrightarrow{\frac{a}{2}C_{n,m}p^mq^{n-m}} (\beta,m),\quad (\alpha,\beta)\in D_1\setminus D_2,\\
(\alpha,n) &\xrightarrow{\frac{a}{2}C_{n,m}q^mp^{n-m}} (\beta,m),\quad (\alpha,\beta)\in D_2\setminus D_1,\\
(\alpha,n) &\xrightarrow{\frac{a}{2}C_{n,m}(p^mq^{n-m}+q^mp^{n-m})} (\beta,m),\quad (\alpha,\beta)\in D_1\cap D_2,
\end{aligned}$$

where $C_{n,m} = n!/m!(n-m)!$ is the combinatorial number and $p$ is the probability of asymmetric binomial partitioning. Let $p_{\alpha,n}$ denote the probability of being in microstate $(\alpha, n)$. Then the evolution of stochastic gene expression dynamics is governed by the following master equation:

$$\begin{aligned}\dot{p}_{\alpha,n} = &\sum_{m=0}^{n-1} \rho_\alpha \mu_{n-m} p_{\alpha,m} + (n+1) d p_{\alpha,n+1} + \sum_{\substack{\beta\neq\alpha \\ (\beta,\alpha)\notin D_1\cup D_2}} q_{\beta\alpha} p_{\beta,n} \\ &+ \frac{a}{2} \sum_{\substack{\beta\neq\alpha \\ (\beta,\alpha)\in D_1}} \sum_{m=n}^{\infty} C_{m,n} p^n q^{m-n} p_{\beta,m} + \frac{a}{2} \sum_{\substack{\beta\neq\alpha \\ (\beta,\alpha)\in D_2}} \sum_{m=n}^{\infty} C_{m,n} q^n p^{m-n} p_{\beta,m} \\ &- \Big(\rho_\alpha \sum_{m=1}^{\infty} \mu_m + nd + \sum_{\beta\neq\alpha} q_{\alpha\beta}\Big) p_{\alpha,n},\end{aligned} \tag{2}$$

where $\rho_\alpha$ is the total burst production rate (for the mother copy or for the two daughter copies) in cellular state $\alpha$, $\mu_n$ is the burst size distribution of the gene product, $d$ is the degradation rate of the gene product, and $q_{\alpha\beta}$ is the transition rate from cellular state $\alpha$ to cellular state $\beta$. Since there are only two gene states, there are only five possible values for $\rho_\alpha$, which are given by

$$\rho_\alpha = \begin{cases} \rho_0, & \alpha = (r, 0), \\ \rho_1, & \alpha = (r, 1), \\ 2\rho_0, & \alpha = (r, 0, 0), \\ 2\rho_1, & \alpha = (r, 1, 1), \\ \rho_0 + \rho_1, & \alpha = (r, 0, 1), (r, 1, 0), \end{cases}$$

where $\rho_0$ and $\rho_1$ are the burst product rate (for a single gene copy) in the inactive and active gene states, respectively.

### Section S2. Analytical solutions for the autocorrelation function and the power spectrum

To obtain the analytical expression of the autocorrelation function, we first calculate the steady-state factorial moments for gene product abundances. For each cellular state $\alpha$, we introduce the generating function

$$F_\alpha(z) = \sum_{n=0}^{\infty} p_{\alpha,n} z^n.$$

Then Eq. (2) can be converted into the following partial differential equation:

$$\begin{aligned}\partial_t F_\alpha(z) = &\rho_\alpha (H(z) - H(1)) F_\alpha(z) + d(1-z) F_\alpha(z) + \sum_{\beta:(\beta,\alpha)\notin D_1\cup D_2} q_{\beta\alpha} F_\beta(z) \\ &+ \frac{a}{2} \sum_{\beta:(\beta,\alpha)\in D_1} F_\beta(pz+q) + \frac{a}{2} \sum_{\beta:(\beta,\alpha)\in D_2} F_\beta(qz+p),\end{aligned} \tag{3}$$

where

$$H(z) = \sum_{n=1}^{\infty} \mu_n z^n$$

is the generating function of the burst size distribution $\mu_n$. To simplify notation, we define

$$q_{\alpha\beta}^{(1)} = q_{\alpha\beta} I_{(\alpha,\beta)\notin D_1\cup D_2}, \quad q_{\alpha\beta}^{(2)} = \frac{a}{2} I_{(\alpha,\beta)\in D_1}, \quad q_{\alpha\beta}^{(3)} = \frac{a}{2} I_{(\alpha,\beta)\in D_2}. \tag{4}$$

Then Eq. (3) can be simplified as

$$\begin{aligned}\partial_t F_\alpha(z) &= \rho_\alpha(H(z)-H(1))F_\alpha(z) + d(1-z)F_\alpha(z) + \sum_\beta q^{(1)}_{\beta\alpha}F_\beta(z)\\ &\quad + \sum_\beta q^{(2)}_{\beta\alpha}F_\beta(pz+q) + \sum_\beta q^{(3)}_{\beta\alpha}F_\beta(qz+p).\end{aligned} \tag{5}$$

Recall that the (unnormalized) $k$th factorial moment for gene product abundances in cellular state $\alpha$ is defined as

$$m_{k\alpha} := \sum_{n=0}^\infty n(n-1)\cdots(n-k+1)p_{\alpha,n} = F^{(k)}_\alpha(1),$$

where $F^{(k)}_\alpha(1)$ is the $k$th derivative of the generating function $F(z)$ taken value at $z=1$. Taking the $k$th derivative on both sides of Eq. (5), we obtain

$$\begin{aligned}\dot{m}_{k\alpha} &= \sum_{l=0}^{k-1} C_{k,l}H^{(k-l)}(1)\rho_\alpha m_{l\alpha} - kdm_{k\alpha}\\ &\quad + \sum_\beta q^{(1)}_{\beta\alpha}m_{k\beta} + \sum_\beta q^{(2)}_{\beta\alpha}p^k m_{k\beta} + \sum_\beta q^{(3)}_{\beta\alpha}q^k m_{k\beta},\end{aligned} \tag{6}$$

where $C_{k,l} = k!/l!(k-l)!$ is the combinatorial number. To proceed, let $m_k = (m_{k\alpha})$ be a row vector whose components are the $k$th factorial moments in all cellular states and let

$$Q^{(1)} = (q^{(1)}_{\alpha\beta}),\quad Q^{(2)} = (q^{(2)}_{\alpha\beta}),\quad Q^{(3)} = (q^{(3)}_{\alpha\beta})$$

be three square matrices. Then Eq. (6) can be rewritten in the vector form as

$$\dot{m}_k = m_k W_{kk} + \sum_{l=0}^{k-1} m_l W_{lk}, \tag{7}$$

where the matrices $W_{kk}$ and $W_{lk}$ are defined as

$$\begin{aligned}W_{kk} &= Q^{(1)} + p^k Q^{(2)} + q^k Q^{(3)} - kdI,\\ W_{lk} &= C_{k,l}H^{(k-l)}(1)S,\quad l<k,\end{aligned} \tag{8}$$

and $S = \mathrm{diag}(\rho_\alpha)$ is the diagonal matrix whose diagonal entries are the transcription rates in all cellular states. In the non-bursty case, these derivative terms are given by $H'(1) = 1$ and $H^{(k)}(1) = 0$ for any $k \geq 2$; in the bursty case, they are given by $H^{(k)}(1) = k!B^k$, where $B$ is the mean burst size. In particular, when $k=0$, we have

$$\dot{m}_0 = m_0(Q^{(1)} + Q^{(2)} + Q^{(3)}) = m_0 Q.$$

Combining Eqs. (4) and (8), we can see that $W_{kk}$ is a matrix obtained from $Q$ by replacing the rates of transitions in $D_1 \cap D_2$ by $(p^k+q^k)a/2$, replacing the rates of transitions in $D_1 \setminus D_2$ by $p^k a/2$, replacing the rates of transitions in $D_2 \setminus D_1$ by $q^k a/2$, and subtracting $kd$ from the diagonal entries. At the steady state, it follows from Eq. (7) that

$$m_k = -\sum_{l=0}^{k-1} m_l W_{lk} W_{kk}^{-1}. \tag{9}$$

Note that this equation is recursive with respect to $k$. When $k = 0$, $m_0 = \pi = (\pi_\alpha)$ is the steady-state distribution of cellular state transitions with generator matrix $Q$. When $k \geq 1$, $m_k$ can be determined by using the recursive relation.

Let $\alpha(t)$ denote the state of the gene and let $n(t)$ denote the copy number of the gene product in an individual cell at time $t$, respectively. It is then easy to see that

$$\begin{aligned}
\pi_\alpha(t) &= \sum_{n=0}^{\infty} p_{\alpha,n} = \mathbb{P}(\alpha(t) = \alpha), \\
m_{1\alpha}(t) &= \sum_{n=0}^{\infty} n p_{\alpha,n} = \mathbb{E}n(t) I_{\{\alpha(t)=\alpha\}}, \\
m_{2\alpha}(t) &= \sum_{n=0}^{\infty} n(n-1) p_{\alpha,n} = \mathbb{E}n(t)(n(t)-1) I_{\{\alpha(t)=\alpha\}},
\end{aligned}$$

where $m_0 = \pi = (\pi_\alpha)$, $m_1 = (m_{1\alpha})$, and $m_2 = (m_{2\alpha})$ are the first three factorial moments defined recursively in Eq. (9) and $I_A$ is the indicator function of the set $A$. It then follows from Eq. (7) that

$$\begin{aligned}
\dot{\pi}(t) &= \pi(t)Q, \\
\dot{m}_1(t) &= m_1(t)W_{11} + \pi(t)W_{01}, \\
\dot{m}_2(t) &= m_2(t)W_{22} + m_1(t)W_{12} + \pi(t)W_{02},
\end{aligned} \tag{10}$$

where

$$\begin{aligned}
W_{11} &= Q^{(1)} + pQ^{(2)} + qQ^{(3)} - dI, \quad W_{01} = BS, \\
W_{22} &= Q^{(1)} + p^2Q^{(2)} + q^2Q^{(3)} - 2dI, \quad W_{12} = 2BS, \quad W_{02} = H''(1)S.
\end{aligned}$$

Here we have used the fact that $H'(1) = B$ with $B$ being the mean burst size. At the steady state, the first and second factorial moments are given by

$$\begin{aligned}
m_1 &= -\pi W_{01} W_{11}^{-1} = -B\pi S W_{11}^{-1}, \\
m_2 &= -(m_1 W_{12} + \pi W_{02}) W_{22}^{-1} = -(2Bm_1 + H''(1)m_0) S W_{22}^{-1}.
\end{aligned} \tag{11}$$

In the non-bursty case, it is easy to check that $H''(1) = 0$, while in the bursty case, it is easy to check that $H''(1) = 2B^2$, where $B$ is the mean burst size. Since Eq. (10) is a set of linear ordinary differential equations, its time-dependent solution is given by

$$m_1(t) = m_1(0)e^{W_{11}t} + B\int_0^t \pi(0)e^{Qs}Se^{W_{11}(t-s)}\mathrm{d}s. \tag{12}$$

From now on, we assume that the system has reached the steady state. Given the initial state $\alpha(0) = \alpha$ and initial copy number $n(0) = n$ of the gene product, it follows from Eq. (12) that

$$\begin{aligned}
\mathbb{E}n(t) = m_1(t)\mathbb{1} &= m_1(0)e^{W_{11}t}\mathbb{1} + B\int_0^t \pi(0)e^{Qs}Se^{W_{11}(t-s)}\mathbb{1}\mathrm{d}s \\
&= ne_\alpha e^{W_{11}t}\mathbb{1} + B\int_0^t e_\alpha e^{Qs}Se^{W_{11}(t-s)}\mathbb{1}\mathrm{d}s,
\end{aligned}$$

where $e_\alpha$ denotes the row vector whose $\alpha$th component is 1 and all other components are 0. This clearly shows that

$$\mathbb{E}[n(t)|\alpha(0), n(0)] = n(0)e_{\alpha(0)}e^{W_{11}t}\mathbb{1} + B\int_0^t e_{\alpha(0)}e^{Qs}Se^{W_{11}(t-s)}\mathbb{1}\mathrm{d}s.$$

Therefore, at the steady state, we have

$$\begin{aligned}\mathbb{E}n(0)n(t) &= \sum_{\alpha}\mathbb{E}n(0)I_{\{\alpha(0)=\alpha\}}\mathbb{E}[n(t)|\alpha(0),n(0)] \\ &= \sum_{\alpha}\mathbb{E}n(0)I_{\{\alpha(0)=\alpha\}}[n(0)e_{\alpha(0)}e^{W_{11}t}\mathbb{1}+B\int_0^t e_{\alpha(0)}e^{Qs}Se^{W_{11}(t-s)}\mathbb{1}\mathrm{d}s] \\ &= \sum_{\alpha}\mathbb{E}n(0)^2I_{\{\alpha(0)=\alpha\}}e_{\alpha}e^{W_{11}t}\mathbb{1}+B\int_0^t\sum_{\alpha}\mathbb{E}n(0)^2I_{\{\alpha(0)=\alpha\}}e_{\alpha}e^{Qs}Se^{W_{11}(t-s)}\mathbb{1}\mathrm{d}s \\ &= \sum_{\alpha}(m_{1\alpha}+m_{2\alpha})e_{\alpha}e^{W_{11}t}\mathbb{1}+B\int_0^t m_{1\alpha}e_{\alpha}e^{Qs}Se^{W_{11}(t-s)}\mathbb{1}\mathrm{d}s \\ &= (m_1+m_2)e^{W_{11}t}\mathbb{1}+B\int_0^t m_1e^{Qs}Se^{W_{11}(t-s)}\mathbb{1}\mathrm{d}s,\end{aligned}$$

where $m_1 = (m_{1\alpha})$ and $m_2 = (m_{2\alpha})$ are the steady-state factorial moments given in Eq. (11). Since the autocorrelation function is defined as $R(t) = \mathbb{E}n(0)n(t) - \mathbb{E}n(0)\mathbb{E}n(t)$, we finally obtain the explicit expression of the autocorrelation function, which is given by

$$R(t) = (m_1+m_2)e^{W_{11}t}\mathbb{1} - \langle n\rangle^2 + B\int_0^t m_1e^{Qs}Se^{W_{11}(t-s)}\mathbb{1}\mathrm{d}s. \tag{13}$$

This is exactly Eq. (1) in the main text, where $W_{11}$ is abbreviated as $W$. In the regime of fast promoter switching, i.e. $u, v, f \ll d$, we have $W_{11} \approx -dI$ and thus the autocorrelation function can be simplified as

$$R(t) = \langle n^2\rangle e^{-dt} - \langle n\rangle^2 + B\int_0^t m_1e^{Qs}S\mathbb{1}e^{d(s-t)}\mathrm{d}s,$$

where we have used the fact that $\langle n^2\rangle = (m_1+m_2)\mathbb{1}$.

So far, we have obtained the analytical expression of the autocorrelation function $R(t)$ in matrix form. We next write it more explicitly in terms of simple functions. Without loss of generality, we assume that all eigenvalues of $Q$, as well as all eigenvalues of $W_{11}$, are mutually distinct (in fact, any matrix can be approximated by such matrices to any degree of accuracy). Let $\lambda_0\cdots,\lambda_{K-1}$ be all eigenvalues of $Q$ and let $\lambda_K,\cdots,\gamma_{2K-1}$ be all eigenvalues of $W_{11}$. By the Perron-Frobenius theorem, one eigenvalue of the generator matrix $Q$ must be zero and other eigenvalues must have negative real parts. Similarly, all eigenvalues of $W_{11}$ must have negative real parts, which implies that

$$\lambda_0 = 0, \quad \mathrm{Re}(\lambda_1),\cdots,\mathrm{Re}(\lambda_{2K-1}) < 0,$$

where $\mathrm{Re}(x)$ denotes the real part of $x$. From Eq. (13), it can be deduced that the autocorrelation function is a linear combination of exponential functions:

$$R(t) = \sum_{k=1}^{2K-1} u_ke^{\lambda_kt}, \tag{14}$$

where $u_k$ are suitable constants. Taking the derivatives on both sides of Eqs. (13) and (14) and evaluating at $t = 0$ yield

$$\sum_{k=1}^{2K-1} u_k\lambda_k^n = v_n,$$

where

$$v_n = (m_1+m_2)W_{11}^n\mathbb{1} + B\sum_{k=1}^{n} m_1Q^{n-k}SW_{11}^{k-1}\mathbb{1}. \tag{15}$$

This equation can be rewritten in matrix form as

$$\begin{pmatrix} \lambda_1 & \lambda_2 & \cdots & \lambda_{2K-1} \\ \lambda_1^2 & \lambda_2^2 & \cdots & \lambda_{2K-1}^2 \\ \vdots & \vdots & \ddots & \vdots \\ \lambda_1^{2K-1} & \lambda_2^{2K-1} & \cdots & \lambda_{2K-1}^{2K-1} \end{pmatrix} \begin{pmatrix} u_1 \\ u_2 \\ \vdots \\ u_{2K-1} \end{pmatrix} = \begin{pmatrix} v_1 \\ v_2 \\ \vdots \\ v_{2K-1} \end{pmatrix},$$

where the matrix $V$ on the left-hand side is a Vandermonde matrix. This shows that

$$\begin{pmatrix} u_1 \\ u_2 \\ \vdots \\ u_{2K-1} \end{pmatrix} = \begin{pmatrix} \lambda_1 & \lambda_2 & \cdots & \lambda_{2K-1} \\ \lambda_1^2 & \lambda_2^2 & \cdots & \lambda_{2K-1}^2 \\ \vdots & \vdots & \ddots & \vdots \\ \lambda_1^{2K-1} & \lambda_2^{2K-1} & \cdots & \lambda_{2K-1}^{2K-1} \end{pmatrix}^{-1} \begin{pmatrix} v_1 \\ v_2 \\ \vdots \\ v_{2K-1} \end{pmatrix}.$$

To proceed, recall that the inverse of the Vandermonde matrix is given by $V^{-1} = (b_{kl})$, where

$$b_{kl} = (-1)^{l-1} \left[ \frac{\prod\limits_{\substack{1\leq m_1<\cdots m_{2K-1-l}\leq 2K-1 \\ m_1,\cdots,m_{2K-1}\neq k}} \lambda_{m_1}\cdots\lambda_{m_{2K-1-l}}}{\lambda_k \prod\limits_{\substack{1\leq m\leq 2K-1 \\ m\neq k}} (\lambda_m - \lambda_k)} \right]. \tag{16}$$

Therefore, the coefficients $u_k$ are given by

$$u_k = \sum_{l=1}^{2K-1} b_{kl} v_l.$$

Finally, the autocorrelation function can be written as

$$R(t) = \sum_{k=1}^{2K-1} u_k e^{\lambda_k t} = \sum_{k,l=1}^{2K-1} b_{kl} v_l e^{\lambda_k t},$$

where $b_{kl}$ is given in Eq. (16) and $v_l$ is given in Eq. (15). In particular, the autocorrelation function is a linear combination of $2K-1$ exponentially functions. Taking the Fourier transform of the autocorrelation function gives the power spectrum

$$G(\xi) = -\sum_{k=2}^{2K} \frac{2u_k\lambda_k}{4\pi^2\xi^2 + \lambda_k^2}, \tag{17}$$

which is linear combination of $2K-1$ Lorentzian functions. This is exactly Eq. (2) in the main text.

In the regime of fast promoter switching, i.e. $d, f \ll u, v$, gene switching dynamics will reach rapid equilibrium and thus the two gene states can be combined into a single one. In this case, the cellular state is only determined by the $N$ cell cycle stages and $Q$ and $W$ reduce to $N \times N$ matrices. Specifically, the generator matrix $Q$ for cellular state transitions is given by

$$Q = \begin{pmatrix} -a & a & & & \\ & -a & a & & \\ & & \ddots & \ddots & \\ & & & -a & a \\ a & & & & -a \end{pmatrix},$$

and the matrix $W$ is given by

$$W = \begin{pmatrix} -a & a & & & \\ & -a & a & & \\ & & \ddots & \ddots & \\ & & & -a & a \\ a/2 & & & & -a \end{pmatrix}.$$

The eigenvalues of $Q$ can be calculated explicitly as

$$\lambda_k = -a + a\omega_k, \quad 0 \leq k \leq N-1, \tag{18}$$

and the eigenvalues of $W$ can be calculated explicitly as

$$\lambda_{N+k} = -d - a + 2^{-1/N} a\omega_k, \quad 0 \leq k \leq N-1, \tag{19}$$

where $\omega_k = \exp(2k\pi/N)$ are all the $N$th roots of unity. Note that $Q$ is a circulant matrix. It is easy to check that there exists a complex orthogonal matrix

$$R = \frac{1}{\sqrt{N}} \begin{pmatrix} 1 & 1 & \cdots & 1 \\ \omega_0 & \omega_1 & \cdots & \omega_{N-1} \\ \cdots & \cdots & \cdots & \cdots \\ \omega_0^{N-1} & \omega_1^{N-1} & \cdots & \omega_{N-1}^{N-1}, \end{pmatrix}$$

such that $Q$ is diagonalized, i.e.

$$\bar{R}'QR = \begin{pmatrix} \lambda_0 & & & \\ & \lambda_1 & & \\ & & \ddots & \\ & & & \lambda_{N-1} \end{pmatrix}$$

where $\bar{R}'$ is the conjugate transpose of $R$. Similarly, $W$ can be diagonalized as

$$\bar{R}'M^{-1}WMR = \begin{pmatrix} \lambda_N & & & \\ & \lambda_{N+1} & & \\ & & \ddots & \\ & & & \lambda_{2N-1} \end{pmatrix},$$

where $M$ is a diagonal matrix which is given by

$$M = \begin{pmatrix} 1 & & & \\ & 2^{-1/N} & & \\ & & \ddots & \\ & & & 2^{-(N-1)/N} \end{pmatrix}.$$

With these notation, it follows from Eq. (13) that the coefficients $u_k, 0 \leq k \leq N-1$ associated with the eigenvalues of $Q$ are given by

$$\begin{aligned} u_k = {} & [(m_1 + m_2)MR]_k [\bar{R}'M^{-1}\mathbb{1}]_k \\ & - B \sum_{i=1}^{N} \frac{[m_1 R]_i [\bar{R}'SMR]_{ik} [\bar{R}'M^{-1}\mathbb{1}]_k}{\lambda_{i-1} - \lambda_{N+k-1}}, \end{aligned} \tag{20}$$

and the coefficients $u_k, N \leq k \leq 2N-1$ associated with the eigenvalues of $W$ are given by

$$u_k = B\sum_{j=1}^{N} \frac{[m_1R]_k[\bar{R}'SMR]_{kj}[\bar{R}'M^{-1}\mathbb{1}]_j}{\lambda_{k-1}-\lambda_{N+j-1}}. \tag{21}$$

Note that if the cell cycle frequency is increased from $f$ to $\alpha f$ with some $\alpha > 1$, while keeping $\eta = d/f$ and $\langle n\rangle$ as fixed, then it follows from Eqs. (18) and (19) that all the eigenvalues $\lambda_k, 0 \leq k \leq 2N-1$ will become $\alpha\lambda_k$. In addition, it follows from Eq. (11) that both $m_1$ and $m_2$ are invariant under the above transformation. Since $S$ is replaced by $\alpha S$ under the above transformation, it follows from Eqs. (20) and (21) that all the coefficients $u_k, 0 \leq k \leq 2N-1$ will remain the same. These results show that the power spectrum (autocorrelation function) will be stretched (compressed) along the horizontal axis by a factor of $\alpha$.

### Section S3. Analytical expressions for the gene expression mean

Since $m_{1\alpha} = \sum_n np_{\alpha,n}$, the steady-state mean of the gene product abundance is given by

$$\langle n\rangle = m_1\mathbb{1} = -B\pi SW^{-1}\mathbb{1}.$$

In the regime of fast switching, i.e. $f, d \ll u, v$, straightforward calculations show that

$$\langle n\rangle = \frac{\rho_{\text{eff}}B}{\eta f}\left[w+\kappa(1-w)+\frac{(\kappa-1)a_{\eta,N}^w+1-\kappa a_{\eta,N}}{\eta(2a_{\eta,N}-1)}\right], \tag{22}$$

where $w = N_0/N$ is the proportion of cell cycle before gene replication, $\eta = d/f$ is the ratio of the degradation rate and the cell cycle frequency, and $a_{\eta,N} = (1+\eta/N)^N \approx e^\eta$ when $N \gg 1$. This is exactly Eq. (3) in the main text.

For stable gene products, it follows from Eq. (22) and L'Hôpital's rule that

$$\langle n\rangle = \frac{\rho_{\text{eff}}B}{2f}\left[3\kappa-(\kappa-1)w(4-w)+\frac{1}{N}(\kappa w+1-w)\right]. \tag{23}$$

### Section S4. Simplification of the power spectrum for stable gene products

Here we derive the simplified expression of the power spectrum for stable gene products. We first calculate the coefficients $u_k$ associated with $W$. From Eq. (21), when $\eta \ll 1$, straightforward computations show that

$$u_k = \frac{\rho_{\text{eff}}^2B^2\bar{\omega}_k}{f^2N^4|1-\omega_k|^2}\left|(\kappa-1)\left(2N-N_0+\frac{1-\omega_k^{N_0}}{1-\omega_k}\right)+(2-\kappa)N\right|^2.$$

when $N \gg 1$. When $N$ is large, direct computations show that

$$\sum_{k=N}^{2N-1}\frac{2u_k\lambda_k}{4\pi^2\xi^2+\lambda_k^2} \approx \frac{\rho_{\text{eff}}^2B^2N}{2f}\sum_{k=1}^{[N/2]}H_k(\kappa,w)G_k(\xi), \tag{24}$$

where

$$H_k(\kappa,w) = \left[w+\kappa(1-w)+\frac{(\kappa-1)\sin(2kw\pi)}{2k\pi}\right]^2+\frac{(\kappa-1)^2\sin^4(kw\pi)}{k^2\pi^2}$$

is a function of $\kappa$ and $w$ and

$$G_k(\xi) = \frac{2k^2\pi^2 f^2 - \left(\pi^2\xi^2 - a^2\cos\theta_k \sin^2\theta_k/2\right)}{4k^6\pi^6 f^4 + N^2\left(\pi^2\xi^2 - a^2\cos\theta_k\sin^2\theta_k/2\right)^2}.$$

This is exactly Eq. (6) in the main text. Similarly, when $N$ is large, we can prove that

$$\sum_{k=1}^{N-1} \frac{2u_k\lambda_k}{4\pi^2\xi^2 + \lambda_k^2} \approx \frac{2A\langle n\rangle f}{3(4\pi^2\xi^2 + \lambda_N^2)}, \tag{25}$$

where

$$A = 2(\ln 2)^2 J(\kappa, w)\langle n\rangle \left(\gamma(p) + \frac{2}{3N}\right) + \frac{1}{(\ln 2)^2} + \frac{2B}{3\ln 2}\left(\gamma(p) - 2 + \frac{3}{\ln 2}\right),$$

if gene expression is bursty and

$$A = 2(\ln 2)^2 J(\kappa, w)\langle n\rangle \left(\gamma(p) + \frac{2}{3N}\right) + \frac{1}{(\ln 2)^2},$$

if gene expression is non-bursty, where $\gamma(p) = 2(1-4pq)/(1+2pq)$ is a function of $p$ and

$$J(\kappa, w) = \frac{[w + \kappa(1-w)]^2}{\left[\kappa - \frac{1}{3}(\kappa-1)w(4-w)\right]^2} \tag{26}$$

is a function of $\kappa$ and $w$. It then follows from Eq. (24) and (25) that

$$G(\xi) \approx \frac{2A\langle n\rangle f}{3(4\pi^2\xi^2 + \lambda_N^2)} + \frac{\rho_{\text{eff}}^2 B^2 N}{2f}\sum_{k=1}^{[N/2]} H_k(\kappa, w) G_k(\xi). \tag{27}$$

This is exactly Eq. (5) in the main text. From the above equation, when $N \gg 1$, the absolute height of the zero peak is given by

$$\begin{aligned} H_{\text{zero}} &\approx \frac{2A\langle n\rangle f}{3\lambda_N^2} + \frac{\rho_{\text{eff}}^2 B^2 N}{2f}\sum_{k=1}^{\infty} H_k(\kappa, w) G_k(0) \\ &\approx \frac{2A\langle n\rangle}{3(\ln 2)^2 f} + \frac{2}{9}\langle n\rangle^2 f N \sum_{k=1}^{\infty} J_k(\kappa, w)\frac{3}{N^2k^2\pi^2 f^2} \\ &\approx \frac{2A'\langle n\rangle}{3(\ln 2)^2 f}, \end{aligned}$$

where

$$J_k(\kappa, w) = \frac{H_k(\kappa, w)}{\left[\kappa - \frac{1}{3}(\kappa-1)w(4-w)\right]^2}$$

is a function of $\kappa$ and $w$ and

$$A' = 2(\ln 2)^2\langle n\rangle \left[J(\kappa, w)\left(\gamma(p) + \frac{2}{3N}\right) + \frac{C(\kappa, w)}{2\pi^2 N}\right] + \frac{1}{(\ln 2)^2} + \frac{2B}{3\ln 2}\left(\gamma(p) - 2 + \frac{3}{\ln 2}\right),$$

if gene expression is bursty and

$$A' = 2(\ln 2)^2\langle n\rangle \left[J(\kappa, w)\left(\gamma(p) + \frac{2}{3N}\right) + \frac{C(\kappa, w)}{2\pi^2 N}\right] + \frac{1}{(\ln 2)^2},$$

if gene expression is non-bursty. Moreover, the absolute height of the off-zero spectral peak is given by

$$H_{\text{off-zero}} \approx \frac{2}{9}\langle n\rangle^2 fNJ_1(\kappa,w)\frac{2\pi^2 f^2}{4\pi^6 f^4} = \frac{J_1(\kappa,w)\langle n\rangle^2 N}{9\pi^4 f}.$$

Since we have normalized the power spectrum so that $G(0)=1$, the height of the off-zero peak is then the ratio of the absolute heights of the off-zero and zero peaks, which is given by

$$H = \frac{H_{\text{off-zero}}}{H_{\text{zero}}} \approx \frac{J_1(\kappa,w)\langle n\rangle^2 N}{9\pi^4 f}\cdot\frac{3(\ln 2)^2 f}{2A'\langle n\rangle} = \frac{(\ln 2)^2 J_1(\kappa,w)\langle n\rangle N}{6\pi^4 A'}.$$

This can be written more explicitly as

$$H \approx \frac{J_1(\kappa,w)\langle n\rangle N}{\pi^4\left\{6\langle n\rangle\left[2J(\kappa,w)\left(\gamma(p)+\frac{2}{3N}\right)+\frac{C(\kappa,w)}{\pi^2 N}\right]+\frac{6}{(\ln 2)^4}+\frac{4B}{(\ln 2)^3}\left(\gamma(p)-2+\frac{3}{\ln 2}\right)\right\}},$$

if gene expression is bursty and

$$H \approx \frac{J_1(\kappa,w)\langle n\rangle N}{\pi^4\left\{6\langle n\rangle\left[2J(\kappa,w)\left(\gamma(p)+\frac{2}{3N}\right)+\frac{C(\kappa,w)}{\pi^2 N}\right]+\frac{6}{(\ln 2)^4}\right\}},$$

if gene expression is non-bursty. Finally, note that $3(\log 2)^3 \approx 0.9991 \approx 1$. Using this approximation, we obtain

$$H \approx \frac{J_1(\kappa,w)\langle n\rangle N}{6\pi^4\left\{\langle n\rangle\left[2J(\kappa,w)(\gamma(p)+\frac{2}{3N})+\frac{C(\kappa,w)}{\pi^2 N}\right]+\frac{3}{\ln 2}+2B\left(\gamma(p)-2+\frac{3}{\ln 2}\right)\right\}}, \tag{28}$$

if gene expression is bursty and

$$H \approx \frac{J_1(\kappa,w)\langle n\rangle N}{6\pi^4\left\{\langle n\rangle\left[2J(\kappa,w)(\gamma(p)+\frac{2}{3N})+\frac{C(\kappa,w)}{\pi^2 N}\right]+\frac{3}{\ln 2}\right\}},$$

This is exactly Eq. (7) in the main text.

In the special case of symmetric cell division ($p=0.5$), large gene expression mean ($\langle n\rangle \gg 1$), midway replication ($w=0.5$), and no dosage compensation ($\kappa=2$), by analyzing the monotonicity of the function $G(\xi)$ given in Eq. (17), it can be proved the system has a type I spectrum when $N\le 6$, type II spectrum when $7\le N\le 28$, and type III spectrum when $N\ge 29$. The reason why $N\ge 29$ yields a type III spectrum can also be understood from Eq. (28), from which it is easy to check that when $p=0.5$, $w=0,5$, $\kappa=2$, and $\langle n\rangle \gg 1$, the height of the off-zero peak reduces to

$$H \approx \frac{J_1(\kappa,w)N^2}{2\pi^2[4\pi^2 J(\kappa,w)+3C(\kappa,w)]}$$

where

$$J_1(\kappa,w)=1.1716,\quad J(\kappa,w)=1.1211,\quad C(\kappa,w)=1.8943.$$

Straightforward computations shows that $H\ge 1$ if and only if $N\ge 29$. This again shows that type III spectra occur when $N\ge 29$.

Moreover, in the special case of symmetric cell division, large gene expression mean, and strong dosage compensation ($\kappa=1$), by analyzing the monotonicity of the function $G(\xi)$ given in Eq. (17), it can be proved the system has a type I spectrum when $N\le 6$, type II spectrum when $7\le N\le 29$, and type III spectrum when $N \geq 29$. The reason why $N \geq 30$ yields a type III spectrum can also be understood from Eq. (28), from which it is easy to check that when $p = 0.5$, $\kappa = 1$, and $\langle n \rangle \gg 1$, the height of the off-zero peak is given by

$$H \approx \frac{N^2}{9\pi^4}.$$

Straightforward computations shows that $H \geq 1$ if and only if $N \geq 30$. This again shows that type III spectra occur when $N \geq 30$.

### Section S5. Simplification of the power spectrum for unstable gene products

Here we derive the simplified expression of the power spectrum for unstable gene products. We first calculate the coefficients $u_k$ associated with $W$. From Eq. (21), when $\eta \gg 1$, straightforward computations show that

$$u_k = \frac{(\kappa-1)^2\rho_{\text{eff}}^2 B^2}{N^2\eta^2 f^2}\left|\frac{1-\omega_k^{N_0}}{1-\omega_k}\right|^2.$$

When $N$ is large, direct computations show that

$$\sum_{k=N}^{2N-1}\frac{2u_k\lambda_k}{4\pi^2\xi^2+\lambda_k^2} \approx \frac{2(\kappa-1)^2\rho_{\text{eff}}^2 B^2 N\sin^2(kw\pi)}{\eta^2 f}\sum_{k=1}^{[N/2]}G_k(\xi), \tag{29}$$

where

$$G_k(\xi) = \frac{2k^2\pi^2 f^2 + \left(\pi^2\xi^2 - a^2\cos\theta_k\sin^2\theta_k/2\right)}{4k^6\pi^6 f^4 + N^2\left(\pi^2\xi^2 - a^2\cos\theta_k\sin^2\theta_k/2\right)^2}.$$

Similarly, when $N$ is large, we can prove that

$$\sum_{k=1}^{N-1}\frac{2u_k\lambda_k}{4\pi^2\xi^2+\lambda_k^2} \approx 0. \tag{30}$$

It then follows from Eq. (29) and (30) that

$$G(\xi) \approx \frac{2A\langle n\rangle f}{3(4\pi^2\xi^2+\lambda_N^2)} + \frac{\rho_{\text{eff}}^2 B^2 N}{2f}\sum_{k=1}^{[N/2]}G_k(\xi). \tag{31}$$

This is exactly Eq. (12) in the main text. From the above equation, when $N \gg 1$, the absolute height of the zero peak is given by

$$H_{\text{zero}} \approx \frac{\rho_{\text{eff}}^2 B^2 N}{2f}\sum_{k=1}^{\infty}G_k(0) \approx \frac{2C(\kappa-1)^2\rho_{\text{eff}}^2 B^2}{N\pi^2\eta^2 f^3},$$

and the absolute height of the off-zero spectral peak is given by

$$H_{\text{off-zero}} \approx \frac{\rho_{\text{eff}}^2 B^2 N}{2f}\cdot\frac{2\pi^2 f^2}{4\pi^6 f^4} = \frac{(\kappa-1)^2\rho_{\text{eff}}^2 B^2 N\sin(w\pi)}{\pi^4\eta^2 f^3},$$

where

$$C(w) = \sum_{k=1}^{\infty}\frac{\sin^2(kw\pi)}{k^2}$$

is a function of $w$. Since we have normalized the power spectrum so that $G(0) = 1$, the height of the off-zero peak is then the ratio of the absolute heights of the off-zero and zero peaks, which is given by

$$H = \frac{H_{\text{off-zero}}}{H_{\text{zero}}} \approx \frac{\sin(w\pi)N^2}{2C(w)\pi^2}.$$

This is exactly Eq. (9) in the main text.

### Section S6. Master equation for different cell tracking protocols

For asymmetric cell division, recall that the probability for a protein molecule being allocated to one daughter is $p < 1/2$ and the probability of being allocated to the other is $q = 1 - p$. For a cellular state $(N, i, j)$ in cell cycle stage $N$, suppose that $i$ and $j$ record the gene states of the daughter copies that will be allocated to the smaller and larger daughter cell, respectively. If the smaller daughter is tracked after cell division, the cell will transition from some cellular state $(N, i, j)$ in stage $N$ to the cellular state $(1, i)$ in stage 1. Then the master equation describing the evolution of the cellular state dynamics needs modification. Specifically, for $r = 1$, the master equation given in Eq. (1) should be replaced by

$$\begin{aligned}\dot{\pi}_{1,0} &= a(\pi_{N,0,0} + \pi_{N,0,1} - \pi_{1,0}) + (v\pi_{1,1} - u\pi_{1,0}),\\ \dot{\pi}_{1,1} &= a(\pi_{N,1,0} + \pi_{N,1,1} - \pi_{1,1}) + (u\pi_{1,0} - v\pi_{1,1}),\end{aligned}$$

where the first term corresponds to cell division and the second term corresponds to gene switching of the mother copy. The transition diagrams of cellular states under the inheritance and reset mechanisms are illustrated in fig. S3(a),(b), respectively.

Similarly, if the larger daughter is tracked after cell division, the cell will transition from some cellular state $(N, i, j)$ in stage $N$ to the cellular state $(1, j)$ in stage 1. In this case, for $r = 1$, the master equation given in Eq. (1) should be replaced by

$$\begin{aligned}\dot{\pi}_{1,0} &= a(\pi_{N,0,0} + \pi_{N,1,0} - \pi_{1,0}) + (v\pi_{1,1} - u\pi_{1,0}),\\ \dot{\pi}_{1,1} &= a(\pi_{N,0,1} + \pi_{N,1,1} - \pi_{1,1}) + (u\pi_{1,0} - v\pi_{1,1}),\end{aligned}$$

The transition diagrams of cellular states under the inheritance and reset mechanisms are illustrated in fig. S3(c),(d), respectively.

The microstate of the gene of interest can be represented by an order pair $(\alpha, n)$, where $\alpha$ is the cellular state and $n$ is the copy number of the gene product. At division, the cell will transition from some microstate $(N, i, j, n)$ in cell cycle stage $N$ to another microstate $(1, i, m)$ or $(1, j, m)$ in cell cycle stage 1 with $m \leq n$. To model asymmetric cell division, we need to distinguish the following two types of transitions between cellular states:

$$\begin{aligned}D_1 &= \{(\alpha, \beta) : \alpha = (N, i, j),\ \beta = (1, i)\},\\ D_2 &= \{(\alpha, \beta) : \alpha = (N, i, j),\ \beta = (1, j)\},\end{aligned}$$

where $\alpha$ is a cellular state in cell cycle stage $N$ and $\beta$ is a cellular state in cell cycle stage 1. For any pair of cellular states $(\alpha, \beta)$ in $D_1$ or $D_2$, if the smaller daughter is tracked at cell division, the transition rate from $\alpha$ to $\beta$ is given as follows:

$$\begin{aligned}q_{\alpha\beta} &= a, \quad (\alpha, \beta) \in D_1 \setminus D_2,\\ q_{\alpha\beta} &= 0, \quad (\alpha, \beta) \in D_2 \setminus D_1,\\ q_{\alpha\beta} &= a, \quad (\alpha, \beta) \in D_1 \cap D_2;\end{aligned}$$

if the larger daughter is tracked at cell division, the transition rate from $\alpha$ to $\beta$ is given as follows:

$$\begin{aligned}q_{\alpha\beta} &= 0, \quad (\alpha, \beta) \in D_1 \setminus D_2,\\ q_{\alpha\beta} &= a, \quad (\alpha, \beta) \in D_2 \setminus D_1,\\ q_{\alpha\beta} &= a, \quad (\alpha, \beta) \in D_1 \cap D_2;\end{aligned}$$

The transitions in $D_1 \setminus D_2$, $D_2 \setminus D_1$, and $D_1 \cap D_2$ are marked by the red, green, and orange arrows in fig. S3, respectively. Due to asymmetric division, if the smaller daughter is tracked at division, the transition rate from microstate $(\alpha, n)$ to microstate $(\beta, m)$ is given as follows:

$$(\alpha, n) \xrightarrow{aC_{n,m}p^m q^{n-m}} (\beta, m), \quad (\alpha, \beta) \in D_1.$$

Then the evolution of stochastic gene expression dynamics is governed by the master equation

$$\begin{aligned}\dot{p}_{\alpha,n} = & \sum_{m=0}^{n-1} \rho_\alpha \mu_{n-m} p_{\alpha,m} + (n+1)dp_{\alpha,n+1} + \sum_{\substack{\beta\neq\alpha \\ (\beta,\alpha)\notin D_1\cup D_2}} q_{\beta\alpha} p_{\beta,n} \\ & + a \sum_{\substack{\beta\neq\alpha \\ (\beta,\alpha)\in D_1}} \sum_{m=n}^{\infty} C_{m,n} p^n q^{m-n} p_{\beta,m} \\ & - \Big(\rho_\alpha \sum_{m=1}^{\infty} \mu_m + nd + \sum_{\beta\neq\alpha} q_{\alpha\beta}\Big) p_{\alpha,n}.\end{aligned}$$

Similarly, if the larger daughter is tracked at division, the transition rate from microstate $(\alpha, n)$ to microstate $(\beta, m)$ is given as follows:

$$(\alpha, n) \xrightarrow{aC_{n,m}q^m p^{n-m}} (\beta, m), \quad (\alpha, \beta) \in D_2.$$

Then the evolution of stochastic gene expression dynamics is governed by the master equation

$$\begin{aligned}\dot{p}_{\alpha,n} = & \sum_{m=0}^{n-1} \rho_\alpha \mu_{n-m} p_{\alpha,m} + (n+1)dp_{\alpha,n+1} + \sum_{\substack{\beta\neq\alpha \\ (\beta,\alpha)\notin D_1\cup D_2}} q_{\beta\alpha} p_{\beta,n} \\ & + a \sum_{\substack{\beta\neq\alpha \\ (\beta,\alpha)\in D_2}} \sum_{m=n}^{\infty} C_{m,n} q^n p^{m-n} p_{\beta,m} \\ & - \Big(\rho_\alpha \sum_{m=1}^{\infty} \mu_m + nd + \sum_{\beta\neq\alpha} q_{\alpha\beta}\Big) p_{\alpha,n}.\end{aligned}$$

When the smaller or larger daughter is tracked at division, the autocorrelation function also takes the form as in Eq. (13), where $m_1$ and $m_2$ are the vectors defined in Eq. (11), $Q = (q_{\alpha\beta})$ is the generator matrix for cellular state transitions, and

$$W_{11} = Q^{(1)} + pQ^{(2)} + qQ^{(3)} - dI,$$

with $Q^{(i)} = (q_{\alpha\beta}^{(i)})$, $i = 1, 2, 3$ being three matrices. If the smaller daughter is tracked at division, the three matrices are defined by

$$q_{\alpha\beta}^{(1)} = q_{\alpha\beta} I_{\{(\alpha,\beta)\notin D_1\cup D_2\}}, \quad q_{\alpha\beta}^{(2)} = aI_{\{(\alpha,\beta)\in D_1\}}, \quad q_{\alpha\beta}^{(3)} = 0.$$

In this case, $W_{11}$ is a matrix is obtained from $Q$ by replacing the rates of transitions in $D_1$ by $pa$ and subtracting $d$ from the diagonal entries. If the larger daughter is tracked at division, the three matrices are defined by

$$q_{\alpha\beta}^{(1)} = q_{\alpha\beta} I_{\{(\alpha,\beta)\notin D_1\cup D_2\}}, \quad q_{\alpha\beta}^{(2)} = 0, \quad q_{\alpha\beta}^{(3)} = aI_{\{(\alpha,\beta)\in D_2\}}.$$

In this case, $W_{11}$ is a matrix is obtained from $Q$ by replacing the rates of transitions in $D_2$ by $qa$ and subtracting $d$ from the diagonal entries.

### Section S7. Estimation of the power spectrum for each cell lineage

In the main text, we have estimated the averaging power spectrum over cell lineages by means of the Wiener-Khinchin theorem. This method, while accurate, computes the average of numerous cell lineages and thus cannot be applied to estimate the power spectrum of a single cell lineage. To estimate the latter, we fit the single-cell time course data with the following AR($P$) model (here an AR($P$) model stands for an AR model of order $P$):

$$\phi_0 n_t + \phi_1 n_{t-1} + \phi_2 n_{t-2} + \cdots + \phi_P n_{t-P} = \theta_0 \epsilon_t,$$

where $\epsilon_t$ is a white noise satisfying $\langle \epsilon_t \rangle = 0$ and $\langle \epsilon_s \epsilon_t \rangle = \delta_{s,t}$ with $\delta_{s,t}$ being the Kronecker delta, $\phi_0 = 1$, and $\phi_1, \cdots, \phi_P$ are real coefficients satisfying the condition that all the complex zeros of the polynomial $\Phi(z) = \sum_{k=0}^{P} \phi_k z^k$ are outside the unit circle $\{z \in \mathbb{C} : |z| = 1\}$. The AR model is a standard and the most widely used model in time series analysis [1]. The sample mean of the time course data $n(0), n(1), \cdots, n(M-1)$ can be estimated as

$$\langle n \rangle = \frac{1}{M} \sum_{k=0}^{M-1} n(k),$$

and the sample autocorrelation function can be estimated as

$$R_i = \frac{1}{M} \sum_{k=0}^{M-1-i} (n(k) - \langle n \rangle)(n(k+i) - \langle n \rangle).$$

Then the coefficients $\phi_1, \cdots, \phi_p$ can be estimated by solving the following Yule-Walker equation:

$$\begin{pmatrix} R_0 & R_1 & \cdots & R_{P-1} \\ R_1 & R_0 & \cdots & R_{P-2} \\ \vdots & \vdots & \ddots & \vdots \\ R_{P-1} & R_{P-2} & \cdots & R_0 \end{pmatrix} \begin{pmatrix} \phi_1 \\ \phi_2 \\ \vdots \\ \phi_P \end{pmatrix} = \begin{pmatrix} R_1 \\ R_2 \\ \vdots \\ R_P \end{pmatrix}.$$

and the parameter $\theta_0$ can be estimated as

$$\theta_0^2 = R_0 - \sum_{k=1}^{P} \phi_k R_k.$$

Finally, the order $P$ of the AR($P$) model can be determined by minimizing the following Akaike information criterion (AIC):

$$\mathrm{AIC}(P) = \log \theta_0^2(P) + 2\frac{P}{M},$$

where $\theta_0(P)$ is the estimated $\theta_0$ when $P$ is fixed. Specifically, we choose $1 \leq P \leq m_N$ such that AIC($P$) attains its minimum, where $m_M$ is in practice often chosen as $m_M = O(\sqrt{M})$. Once we have determined the order of the AR model and estimated all the parameters, the sample power spectrum can be estimated as [1]

$$G(\xi) = \frac{1}{2\pi} \cdot \frac{\theta_0^2}{\left|\sum_{k=0}^{P} \phi_k e^{-ik\xi}\right|^2}, \quad -\pi \leq \xi \leq \pi.$$

### Section S8. Estimation of $w$ and $\kappa$ for each cell lineage

For a single cell lineage, let $n(t)$ be the time course data at time $t$ and $T_k$ be the $k$th cell division time. Since the burst production rate increases from $\rho_{\text{eff}}$ to $\kappa\rho_{\text{eff}}$ upon replication, we can fit the data between two division times by the following mean-field approximation:

$$\hat{n}(t+1) - \hat{n}(t) = \begin{cases} \rho_{\text{eff}}, & \text{if } t - T_k \leq w(T_{k+1} - T_k), \\ \kappa\rho_{\text{eff}}, & \text{if } t - T_k > w(T_{k+1} - T_k), \end{cases}$$

where $t$ is an arbitrary time point between two consecutive division times $T_k$ and $T_{k+1}$. In other words, we use a piecewise linear function to approximate the gene expression levels between two cell division events. Since $w$ represents the proportion of cell cycle before replication, the gene replication time between $T_k$ and $T_{k+1}$ is on average given by $T_k + w(T_{k+1} - T_k)$. The first part of the piecewise linear approximation is given by

$$\hat{n}(t) = n(T_k) + \rho_{\text{eff}}(t - T_k),$$

for any $T_k \leq t \leq T_k + w(T_{k+1} - T_k)$ and the second part of the piecewise linear approximation is given by

$$\hat{n}(t) = n(T_k) + \rho_{\text{eff}} w(T_{k+1} - T_k) + \kappa\rho_{\text{eff}}[t - T_k - w(T_{k+1} - T_k)],$$

for any $T_k + w(T_{k+1} - T_k) < t \leq T_{k+1}$, where $n(T_k)$ is the gene expression level at time $T_k$. Then the distance between the time course data $n(t)$ and the piecewise linear approximation $\hat{n}(t)$ is given by

$$D = \sum_{t=0}^{M-1} (n(t) - \hat{n}(t))^2. \tag{32}$$

When $\eta = 0$, it follows from Eq. (23) that the gene expression mean is given by

$$\langle n \rangle = \frac{\rho_{\text{eff}} B}{2f} \left[ 3\kappa - (\kappa - 1)w(4 - w) + \frac{1}{N}(\kappa w + 1 - w) \right].$$

The sample mean of the time course data can be estimated as

$$\langle n \rangle = \frac{1}{M} \sum_{k=0}^{M-1} n_k.$$

Therefore, $\rho_{\text{eff}}$ can be represented using $w$ and $\kappa$ as

$$\rho_{\text{eff}} = \frac{2f\langle n \rangle}{3\kappa - (\kappa - 1)w(4 - w) + \frac{1}{N}(\kappa w + 1 - w)}.$$

Finally, the parameters $w$ and $\kappa$ can be estimated by minimizing the distance $D$ in Eq. (32) using the least square criterion.

### Section S9. Estimation of the fluorescence intensity per protein molecule

Suppose that cell division occurs at a particular time. Let $n_b$ be the protein copy number just before division and let $n_a$ be the protein copy number just after division. Since we have assumed symmetric binomial partitioning at cell division, $n_a$ and $n_b$ are related by the following equality:

$$\mathbb{P}(n_a = m) = \sum_{n=m}^{\infty} C_{n,m} \left( \frac{1}{2} \right)^n \mathbb{P}(n_b = n). \tag{33}$$

Therefore, the means of $n_a$ and $n_b$ are related by

$$\begin{aligned}\langle n_a\rangle &= \sum_{m=0}^{\infty} m\mathbb{P}(n_a = m)\\ &= \sum_{m=0}^{\infty} m\sum_{n=m}^{\infty} C_{n,m}\left(\frac{1}{2}\right)^n \mathbb{P}(n_b = n)\\ &= \sum_{n=0}^{\infty}\left(\frac{1}{2}\right)^n \mathbb{P}(n_b = n)\sum_{m=0}^{n} mC_{n,m}\\ &= \sum_{n=0}^{\infty}\left(\frac{1}{2}\right)^n \mathbb{P}(n_b = n)2^{n-1}n\\ &= \frac{1}{2}\sum_{n=0}^{\infty} n\mathbb{P}(n_b = n) = \frac{1}{2}\langle n_b\rangle,\end{aligned}\tag{34}$$

and the second factorial moments of $n_a$ and $n_b$ are related by

$$\begin{aligned}\langle n_a(n_a-1)\rangle &= \sum_{m=0}^{\infty} m(m-1)\mathbb{P}(n_a = m)\\ &= \sum_{m=0}^{\infty} m(m-1)\sum_{n=m}^{\infty} C_{n,m}\left(\frac{1}{2}\right)^n \mathbb{P}(n_b = n)\\ &= \sum_{n=0}^{\infty}\left(\frac{1}{2}\right)^n \mathbb{P}(n_b = n)\sum_{m=0}^{n} m(m-1)C_{n,m}\\ &= \sum_{n=0}^{\infty}\left(\frac{1}{2}\right)^n \mathbb{P}(n_b = n)2^{n-2}n(n-1)\\ &= \frac{1}{4}\sum_{n=0}^{\infty} n(n-1)\mathbb{P}(n_b = n) = \frac{1}{4}\langle n_b(n_b-1)\rangle.\end{aligned}\tag{35}$$

Let $\mu_b$ be the mean of the fluorescence intensity just before division, let $\mu_a$ be the mean of the fluorescence intensity just after division, and let $\beta$ be the fluorescence intensity per protein molecule. It then following from Eq. (34) that

$$\mu_a = \beta\langle n_a\rangle = \frac{1}{2}\beta\langle n_b\rangle = \frac{1}{2}\mu_b.\tag{36}$$

Let $\sigma_b$ be the standard deviation of the fluorescence intensity just before division and let $\sigma_a$ be the standard deviation of the fluorescence intensity just after division. Combining Eqs. (34) and (35) shows that

$$\begin{aligned}\sigma_a^2 &= \beta^2[\langle n_a(n_a-1)\rangle + \langle n_a\rangle - \langle n_a\rangle^2]\\ &= \beta^2\left[\frac{1}{4}\langle n_a(n_a-1)\rangle + \frac{1}{2}\langle n_b\rangle - \frac{1}{4}\langle n_b\rangle^2\right]\\ &= \frac{1}{4}\sigma_b^2 + \frac{1}{4}\beta\mu_b.\end{aligned}\tag{37}$$

Let $\gamma_b = \sigma_b^2/\mu_b$ be the Fano factor of the fluorescence intensity just before division and let $\gamma_a = \sigma_a^2/\mu_a$ be the Fano factor of the fluorescence intensity just after division. Combining Eqs. (36) and (37) finally shows that

$$\gamma_a = \frac{1}{2}\gamma_b + \frac{1}{2}G.$$

This is exactly Eq. (11) in the main text.

### Section S10. Technical details about Fig. 2

In (a),(b), the model parameters are chosen as $w = 0.5, f = 0.1, d = \eta f, \kappa = 2, p = 0.5, \rho_0 = 0, u = v = 10^6$ and gene expression is assumed to be non-bursty. The parameters $\rho_1$ and $\rho_{\text{eff}}$ are chosen so that $\langle n \rangle = 1000$. The rest of the parameters are chosen as $N = 4, N_0 = 2, \eta = 0$ for the type I spectrum, $N = 16, N_0 = 8, \eta = 0.2$ for the type II spectrum, $N = 16, N_0 = 8, \eta = 1.2$ for the type III spectrum, and $N = 16, N_0 = 8, \eta = 3.6$ for the type IV spectrum. The red and green dashed curves in the insets show the decomposition of the power spectrum into the part with a zero peak and the part with an off-zero peak, respectively (see Eq. (4) in the main text for the decomposition). Specifically, the red dashed curve shows the contribution of the eigenvalues of $W$, which mainly controls the zero peak as well as the decay of the spectrum, and the green dashed curve shows the contribution of the eigenvalues of $Q$, which mainly controls the off-zero peak.

In (c), the model parameters are chosen as $w = 0.5, f = 0.1, d = 0.5, \eta = 4, \kappa = 2, p = 0.5, \rho_0 = 0, u = v = 10^6$ and gene expression is assumed to be non-bursty. The parameters $\rho_1$ and $\rho_{\text{eff}}$ are chosen so that $\langle n \rangle = 150$.

In (d), the model parameters are chosen as $N = 50, N_0 = wN, f = 0.1, d = 0.1, \eta = 1, \kappa = 2, \rho_0 = 0, u = v = 10^6$ and gene expression is assumed to be non-bursty. The parameters $\rho_1$ and $\rho_{\text{eff}}$ are chosen so that $\langle n \rangle = 50$.

In (e), the model parameters are chosen as $N = 20, N_0 = 10, w = 0.5, f = 0.1, d = 0.1, \eta = 1, \kappa = 2, p = 0.5, \rho_0 = 0, u = v = 10^6$. The parameters $\rho_1$ and $\rho_{\text{eff}}$ are varied according to the value of $\langle n \rangle$.

In (f), the model parameters are chosen as $N = 100, N_0 = 50, w = 0.5, f = 0.1, d = \eta = 0, \kappa = 2, p = 0.5, \rho_0 = 0, u = v = 10^6$ and gene expression is assumed to be non-bursty. The parameters $\rho_1$ and $\rho_{\text{eff}}$ are chosen so that $\langle n \rangle = 100$. Inset: Same as (f) but for $d = 10$ and $\eta = 100$.

### Section S11. Technical details about Fig. 3

In (a), the model parameters are chosen as $N = 6, N_0 = 3, w = 0.5, f = 0.1, d = \eta f, p = 0.5, \rho_0 = 0, u = v = 10^6$ and gene expression is assumed to be non-bursty. The parameters $\rho_1$ and $\rho_{\text{eff}}$ are chosen so that $\langle n \rangle = 5 \times 10^3$.

In (b), the model parameters are the same as in (a) except $N = 16$ and $N_0 = 8$.

In (c), the model parameters are the same as in (a) except $N = 32$ and $N_0 = 16$.

In (d), the model parameters are chosen as $N = 20, N_0 = 10, w = 0.5, f = 0.1, d = \eta f, \kappa = 2, p = 0.5, \rho_0 = 0, u = v = 10^6$ and gene expression is assumed to be non-bursty. The parameter $\eta$ is chosen as $\eta = 0$ for stable gene products and $\eta = 50$ for unstable gene products. The parameters $\rho_1$ and $\rho_{\text{eff}}$ are chosen so that $\langle n \rangle = 1500$.

In (e), the model parameters are chosen as $N = 20, N_0 = 20, w = 1, f = 0.1, d = 0.1, \eta = 1, p = 0.5, \rho_1 = 200, \rho_0 = 0$ and gene expression is assumed to be non-bursty.

In (f), the model parameters are chosen as $N = 20, N_0 = 20, w = 1, f = 0.1, d = 0.1, \eta = 1, p = 0.5, \rho_1 = 200, \rho_0 = 40$ and gene expression is assumed to be non-bursty.

In (g), the model parameters are chosen as $N = 20, N_0 = 10, w = 0.5, f = 0.1, d = 0.2, \eta = 2, \kappa = 2, \rho_1 = 540, \rho_0 = 0, \rho_{\text{eff}} = 270, u = v = 10^6$ and gene expression is assumed to be non-bursty.

In (h), the model parameters are chosen as $N = 20, N_0 = 10, w = 0.5, f = 0.1, d = 0.2, \eta = 2, \kappa = 2, p = 0.3, \rho_1 = 540, \rho_0 = 0, \rho_{\text{eff}} = 270, u = v = 10^6$ and gene expression is non-bursty.

| Cell types | $N$ | $w$ | Mean doubling time | Median mRNA half-life | $\eta$ for mRNA | Median protein half-life | $\eta$ for protein |
|---|---|---|---|---|---|---|---|
| *E. coli* (K10 + M9 + 30$^{o}$C) | NA | NA | 73 min [2] | 2.1 min (0.6 - 24.6 min) [2] | 24.1 (2.1 - 84.3) | about 20 h [3] | 0.042 |
| *E. coli* (DF261 + M9 + 30$^{o}$C) | NA | NA | 78 min [2] | 2.8 min (0.8 - 43.5 min) [2] | 19.3 (1.2 - 67.6) | about 20 h [3] | 0.045 |
| *E. coli* (N3433 + M9 + 30$^{o}$C) | NA | NA | 75 min [2] | 3.7 min (1.0 - 44.2 min) [2] | 14.1 (1.2 - 52.0) | about 20 h [3] | 0.043 |
| *E. coli* (SU02 + M9 + 30$^{o}$C) | NA | NA | 75 min [2] | 4.0 min (1.6 - 55.5 min) [2] | 13.0 (0.9 - 32.5) | about 20 h [3] | 0.043 |
| *E. coli* (YHC012 + M9 + 30$^{o}$C) | NA | NA | 84 min [2] | 4.0 min (1.2 - 50.2 min) [2] | 14.6 (1.2 - 48.5) | about 20 h [3] | 0.049 |
| *E. coli* (SH3208 + M9 + 30$^{o}$C) | NA | NA | 81 min [2] | 3.2 min (1.1 - 24.6 min) [2] | 17.5 (2.3 - 51.0) | about 20 h [3] | 0.047 |
| *E. coli* (BZ453 + M9 + 30$^{o}$C) | NA | NA | 84 min [2] | 6.0 min (1.7 - 38.6 min) [2] | 9.7 (1.5 - 34.2) | about 20 h [3] | 0.049 |
| *E. coli* (MG1655 + M9 + 30$^{o}$C) | 7 [4] | NA | 30 min [5] | 6.8 min (0.7 - 34.5 min) [5] | 3.1 (0.6 - 29.7) | about 20 h [3] | 0.017 |
| *E. coli* (JM109 + M9 + 37$^{o}$C) | 5 [6] | NA | 87 min [5] | NA | NA | about 20 h [3] | 0.050 |
| *E. coli* (EJ2848 + M9 + 37$^{o}$C) | 9 [7] | NA | 52 min [5] | NA | NA | about 20 h [3] | 0.030 |
| *E. coli* (MC4100 + LB + 25$^{o}$C) | 14 [8] | NA | 68 min [8] | NA | NA | NA | NA |
| *E. coli* (MC4100 + LB + 27$^{o}$C) | 31 [8] | NA | 53 min [8] | NA | NA | NA | NA |
| *E. coli* (MC4100 + LB + 37$^{o}$C) | 31 [8] | NA | 33 min [8] | NA | NA | NA | NA |
| *E. coli* (K12 + AB + 30$^{o}$C) | NA | 0.16 - 0.52 [9] | 125 min [9] | NA | NA | about 20 h [3] | 0.072 |
| *E. coli* (NCM3416 + M9 + 30$^{o}$C) | NA | NA | 90 min [10] | 5.4 min (1.3 - 31.4 min) [10] | 11.6 (2.0 - 48.0) | about 20 h [3] | 0.052 |
| *E. coli* (NCM3416 + LB + 30$^{o}$C) | NA | NA | 30 min [10] | 4.7 min (1.1 - 24.8 min) [10] | 4.4 (0.8 - 18.9) | about 20 h [3] | 0.017 |
| *M. acetivorans* (WWM82 + MeOH + 37$^{o}$C) | NA | NA | 7.5 h [11] | 1.0 h (4 min - 5.8 h) [11] | 5.2 (0.9 - 78.0) | NA | NA |
| *M. acetivorans* (WWM82 + TMA + 37$^{o}$C) | NA | NA | 8.9 h [11] | 1.1 h (14 min - 8.3 h) [11] | 5.6 (0.7 - 26.4) | NA | NA |
| *M. acetivorans* (WWM82 + acetate + 37$^{o}$C) | NA | NA | 24.6 h [11] | 2.8 h (11 min - 9.9 h) [11] | 6.1 (1.7 - 93.0) | NA | NA |
| *S. cerevisiae* | 31 [12] | 0.16 - 0.43 [13] | 1.5 - 2.5 h [14, 15] | 20 min (3 - $>$ 90 min) [16] | 5.2 ($<$ 1.2 - 34.7) | 8.8 h (8 min - $>$ 150 h) [15] | 0.2 ($<$ 0.01 - 13.0) |
| *S. pombe* | 169 [17] | 0.25 - 0.37 [18] | 1.8 - 4.0 h [15, 19] | 33 min (10 - 96 min) [20] | 5.0 (1.7 - 16.6) | 12 h (36 min - $>$ 150 h) [15] | 0.2 ($<$ 0.02 - 4.6) |
| Mouse Fibroblasts (N1H3T3) | 12 [21] | 0.40 - 0.80 [22] | 27.5 h [23] | 9 h (1.4 - 40 h) [23] | 2.1 (0.5 - 13.6) | 46 h (30 min - $>$ 150 h) [23] | 0.4 ($<$ 0.13 - 38.1) |
| Mouse embryonic stem cells | NA | NA | about 12 h [24] | 7.1 h (0.3 - $>$ 24 h) [25] | 1.2 ($<$ 0.3 - 27.7) | NA | NA |
| Mouse epidermal stem cells | NA | NA | about 200 h [26] | NA | NA | NA | NA |
| Human (HeLa) | 17 [27] | 0.65 - 0.83 [28] | 24.7 h [29] | 9.5 h (0.8 - 130 h) [28] | 1.9 (0.13 - 21.4) | NA | NA |
| Human cancer cells (H1299) | 75 [30] | NA | 22.5 h [30] | NA | NA | 9 h (45 min - 22.5 h) [30] | 1.7 (0.7 - 20.8) |
| Human cancer cells (HepG2) | NA | NA | 24 - 48 h [31] | 10 h (0.1 - 27.2 h) [31] | 3.3 (1.2 - 332.7) | 8.7 h (19 min - 36.7 h) [32] | 3.8 (0.9 - 105.1) |
| Human embryonic stem cells | NA | 0.19 - 0.85 [33] | 15.8 h [33] | NA | NA | NA | NA |

Table 1: Table S1. **Biological values of some important model parameters across different cell types, from prokaryotes to yeast then to higher eukaryotes.** The number $N$ of effective cell cycle stages is determined by fitting the distribution of the doubling time to an Erlang distribution. The lower and upper bounds of $w$ are determined as the relative durations of the $G_1$ phase and the $G_1 + S$ phase, respectively. The bracket in the first column gives the strain, medium, and temperature for prokaryotes or the cell type for eukaryotes. Some more data about mRNA half-lives for other *E. coli* strains can be found in [2, 10]. The brackets in the last two columns give the ranges of the mRNA and protein half-lives for all studied genes in an experiment. The value of $\eta$ is calculated using the mean doubling time $T$ and the half-life $T_H$ via the formula $\eta = (\ln 2)T/T_H$.

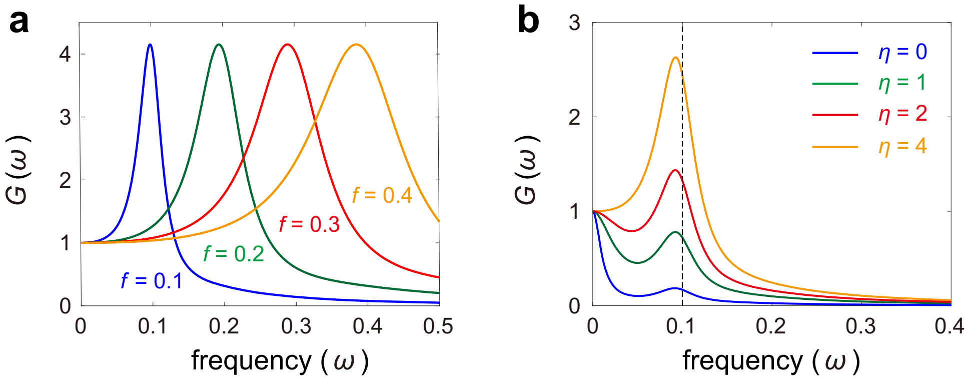


Fig. S1. **Power spectrum and time traces for stochastic gene expression oscillations.** (a) Power spectrum as $f$ varies. Here $f = a/N$ is varied by fixing $N$ and $\langle n \rangle$ and tuning $a$ and $\rho_{\mathrm{eff}}$. If $f$ is increased to $\alpha f$, the power spectrum will be stretched along the frequency axis by a factor of $\alpha$. The model parameters are chosen as $N = 16, N_0 = 8, w = 0.5, \eta = 5, d = \eta f, \kappa = 2, p = 0.5, \rho_0 = 0, u = v = 10^6$ and gene expression is assumed to be non-bursty. The parameters $\rho_1$ and $\rho_{\mathrm{eff}}$ are chosen so that $\langle n \rangle = 150$. (b) Power spectrum as $\eta$ varies, where $\eta = d/f$ is varied by fixing $f$ and $\langle n \rangle$ and tuning $d$ and $\rho_{\mathrm{eff}}$. The model parameters are chosen as $N = 12, N_0 = 6, w = 0.5, f = 0.1, d = \eta f, \kappa = 2, p = 0.5, \rho_0 = 0, u = v = 10^6$ and gene expression is assumed to be non-bursty. The parameters $\rho_1$ and $\rho_{\mathrm{eff}}$ are chosen so that $\langle n \rangle = 150$.

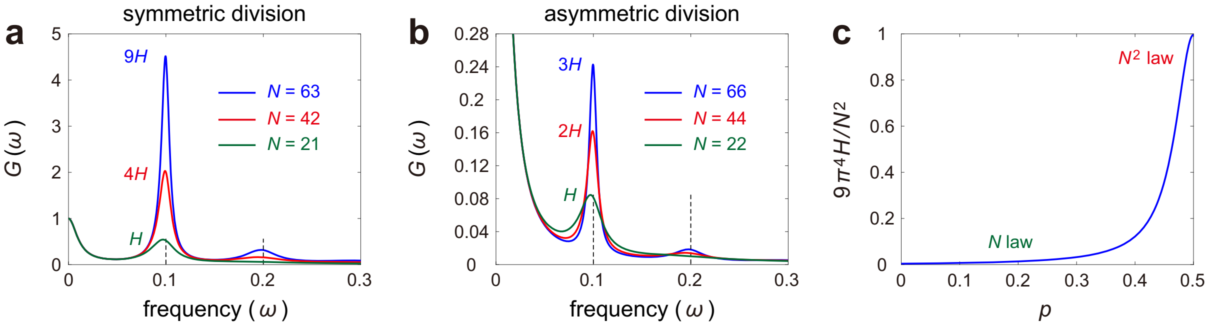


Fig. S2. **Stochastic gene expression oscillations for stable gene products under symmetric and asymmetric cell division.** (a) Power spectrum as $N$ varies under symmetric cell division. The height of the off-zero spectral peak depends on $N$ quadratically when $N$ and $\langle n\rangle$ are both large. The model parameters are chosen as $f = 0.1, d = \eta = 0, \kappa = 1, p = 0.5, \rho_0 = 0, u = v = 10^6$. The parameter $N$ is chosen as $N = 21$ (green curve), $N = 42$ (red curve), and $N = 63$ (blue curve). The parameters $\rho_1$ and $\rho_{\rm eff}$ are chosen so that $\langle n\rangle = 10^5$. (b) Power spectrum as $N$ varies under asymmetric cell division. The height of the off-zero spectral peak depends on $N$ linearly when $N$ and $\langle n\rangle$ are both large. The model parameters are chosen as $f = 0.1, d = \eta = 0, \kappa = 1, p = 0.3, \rho_0 = 0, u = v = 10^6$. The parameter $N$ is chosen as $N = 22$ (green curve), $N = 44$ (red curve), and $N = 66$ (blue curve). The parameters $\rho_1$ and $\rho_{\rm eff}$ are chosen so that $\langle n\rangle = 10^5$. (c) Phase transition of the oscillatory behavior as asymmetry in cell division varies. The figure shows the order parameter $2\pi^2(4\pi^2J + 3C)H/J_1N^2$ versus the control parameter $p$, where $H$ is the height of the off-zero peak. When $N$ and $\langle n\rangle$ are both large, for symmetric division, the height depends on $N$ quadratically as $H \approx J_1N^2/2\pi^2(4\pi^2J + 3C)$, while for asymmetric division, the height depends on $N$ linearly as $H \approx J_1N/12\pi^4J\gamma$. The order parameter approximately equals 1 for symmetric division and is very small for asymmetric division. A sharp transition of the order parameter is observed when the control parameter $p$ deviates from $p = 0.5$. The model parameters are chosen as $N = 100, f = 0.1, d = \eta = 0, \kappa = 1, \rho_0 = 0, u = v = 10^6$ and gene expression is assumed to be non-bursty. The parameters $\rho_1$ and $\rho_{\rm eff}$ are chosen so that $\langle n\rangle = 10^5$.

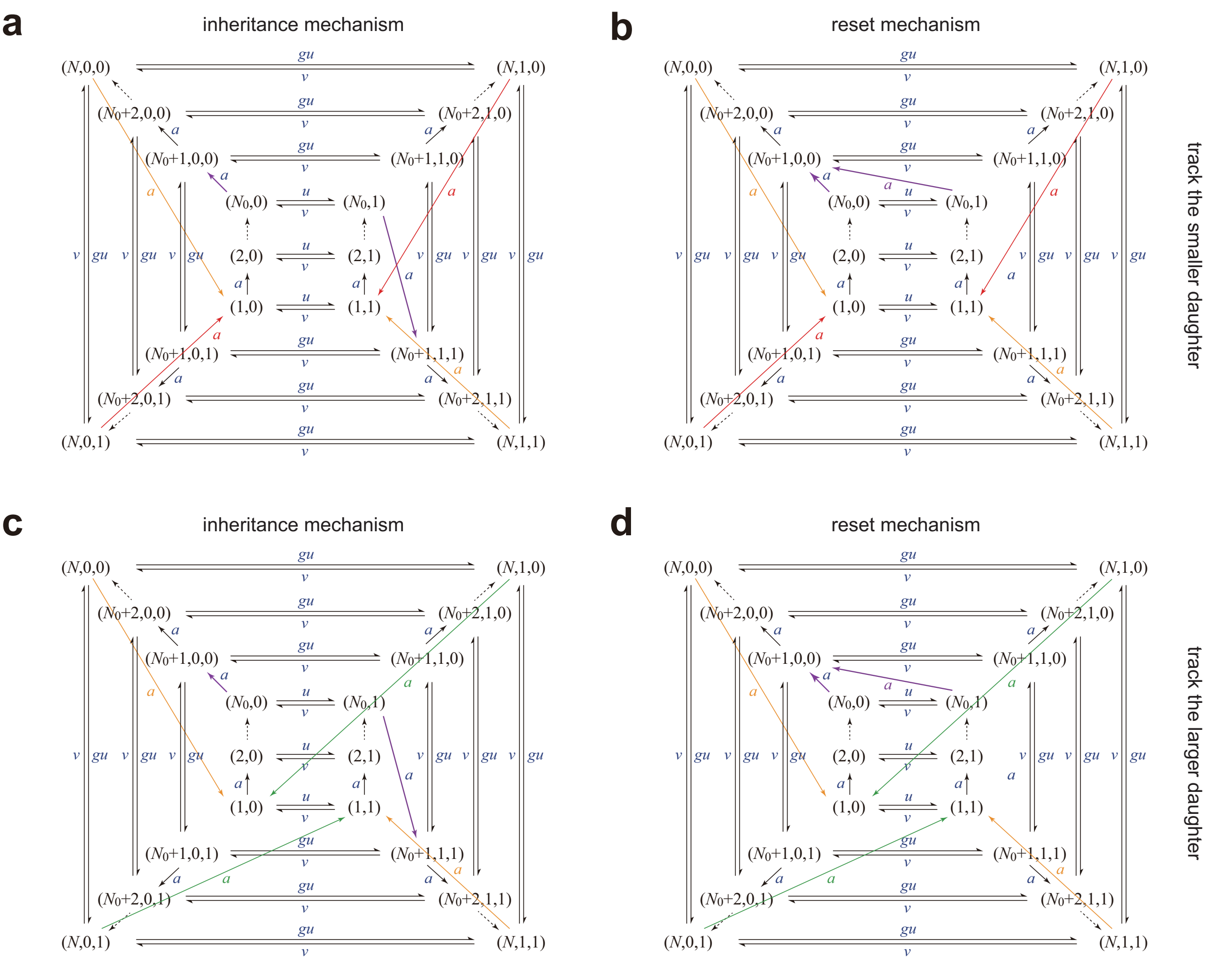


Fig. S3. **Transition diagram of cellular states under two cell tracking protocols.** (a),(b) Transition diagram of cellular states when tracking the smaller daughter after cell division. At division, the smaller daughter with fewer gene products is tracked and thus the cell will transition from cellular state $(N, i, j)$ to cellular state $(1, i)$ (orange and red arrows). (a) Inheritance mechanism. Upon replication, the cell will transition from $(N_0, i)$ to $(N_0, i, i)$ (purple arrows). (b) Reset mechanism. Upon replication, the cell will transition from $(N_0, i)$ to $(N_0, 0, 0)$ (purple arrows). (c),(d) Transition diagram of cellular states when tracking the larger daughter after cell division. At division, the larger daughter with more gene products is tracked and thus the cell will transition from $(N, i, j)$ to $(1, j)$ (orange and green arrows). (c) Inheritance mechanism. (d) Reset mechanism.

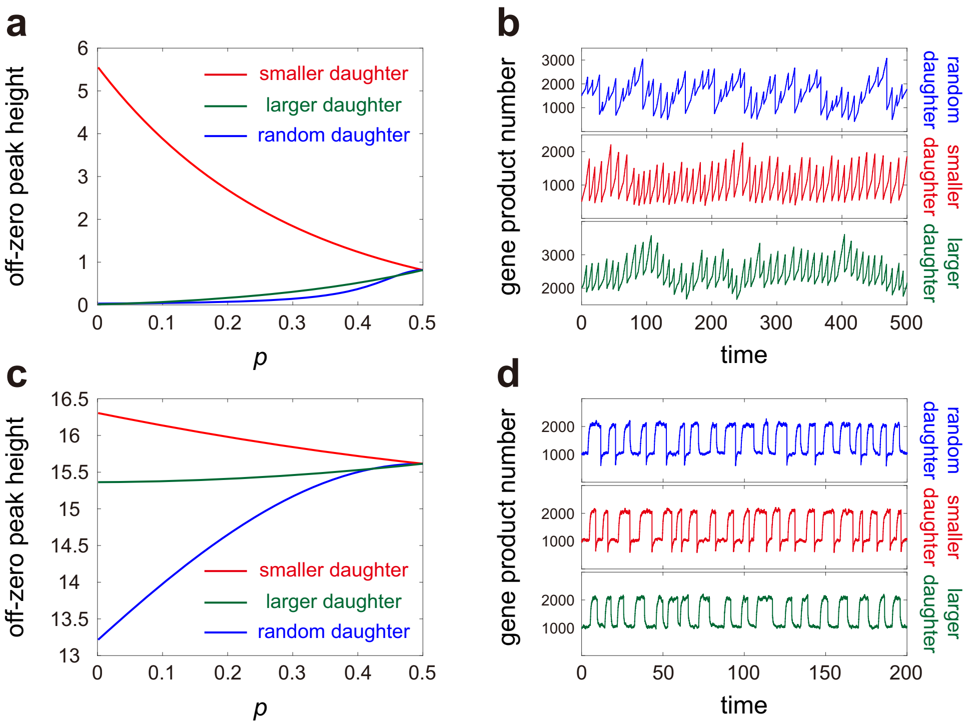


Fig. S4. **Power spectra and stochastic trajectories for three types of tracking protocols.** (a) Height of the off-zero spectral peak (relative to the height of the zero peak) versus $p$ for stable gene products under three types of tracking protocols at cell division: tracking one of the two daughters randomly (blue curve), tracking the smaller daughter (red curve), and tracking the larger daughter (green curve). (b) Typical stochastic trajectories for stable gene products under three types of tracking protocols generated using Gillespie's algorithm. (c) Same as (a) but for unstable gene products. (d) Same as (b) but for unstable gene products. In (a)-(d), the model parameters are chosen as $N = 20, N_0 = 10, w = 0.5, f = 0.1, d = \eta f, \rho_0 = 0, u = v = 10^6$ and gene expression is assumed to be non-bursty. The parameter $\eta$ is chosen as $\eta = 0.2$ in (a),(b) and $\eta = 20$ in (c),(d). The parameter $\rho_1$ and $\rho_{\text{eff}}$ are chosen so that $\langle n \rangle = 150$ for the random tracking protocol. The parameter $p$ is chosen as $p = 0.3$ in (b),(d).

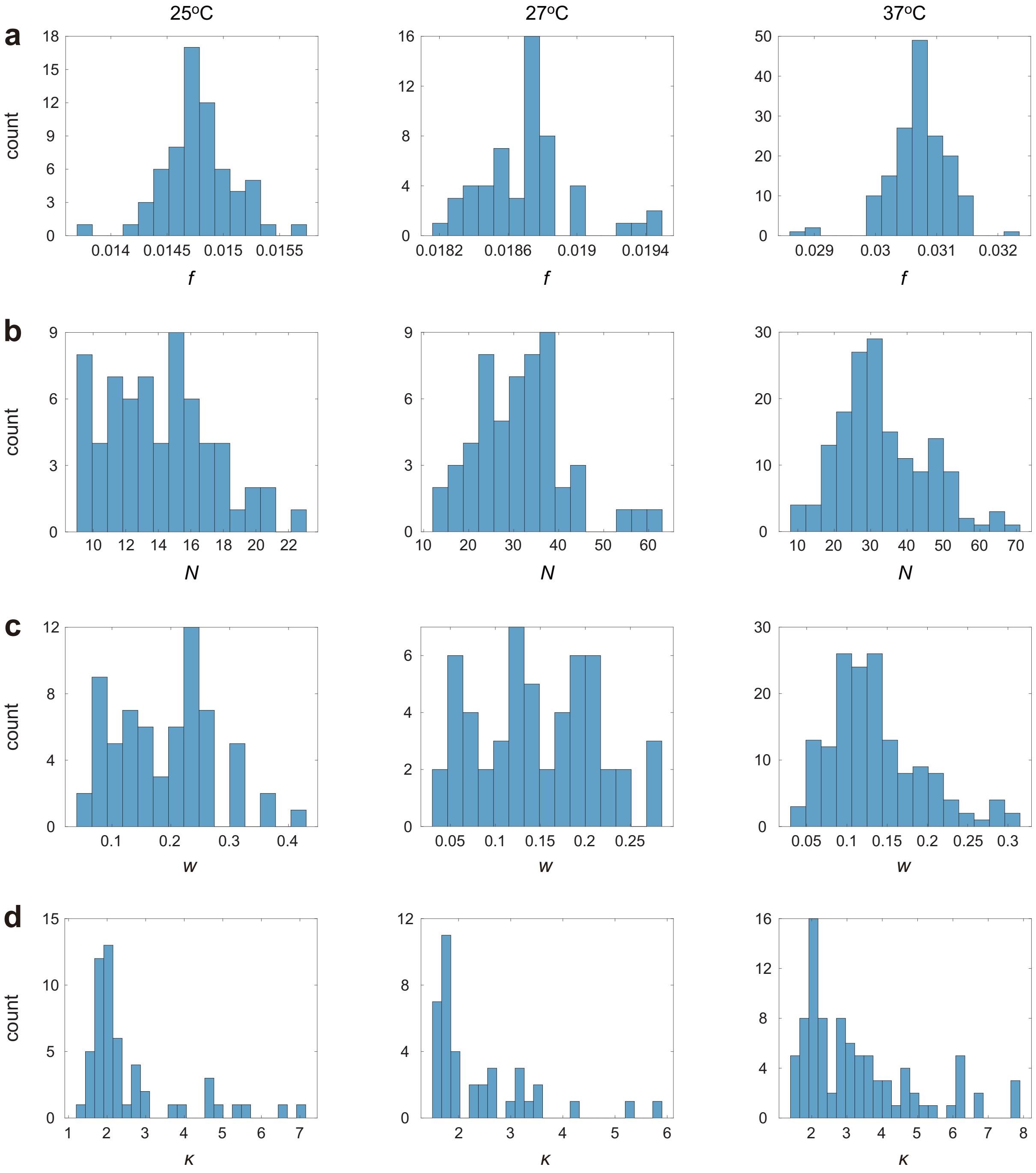


Fig. S5. **Distribution of the first set of model parameters at the three temperatures.** (a) Distribution of the estimated $f$. (b) Distribution of the estimated $N$. (c) Distribution of the estimated $w$. (d) Distribution of the estimated $\kappa$.

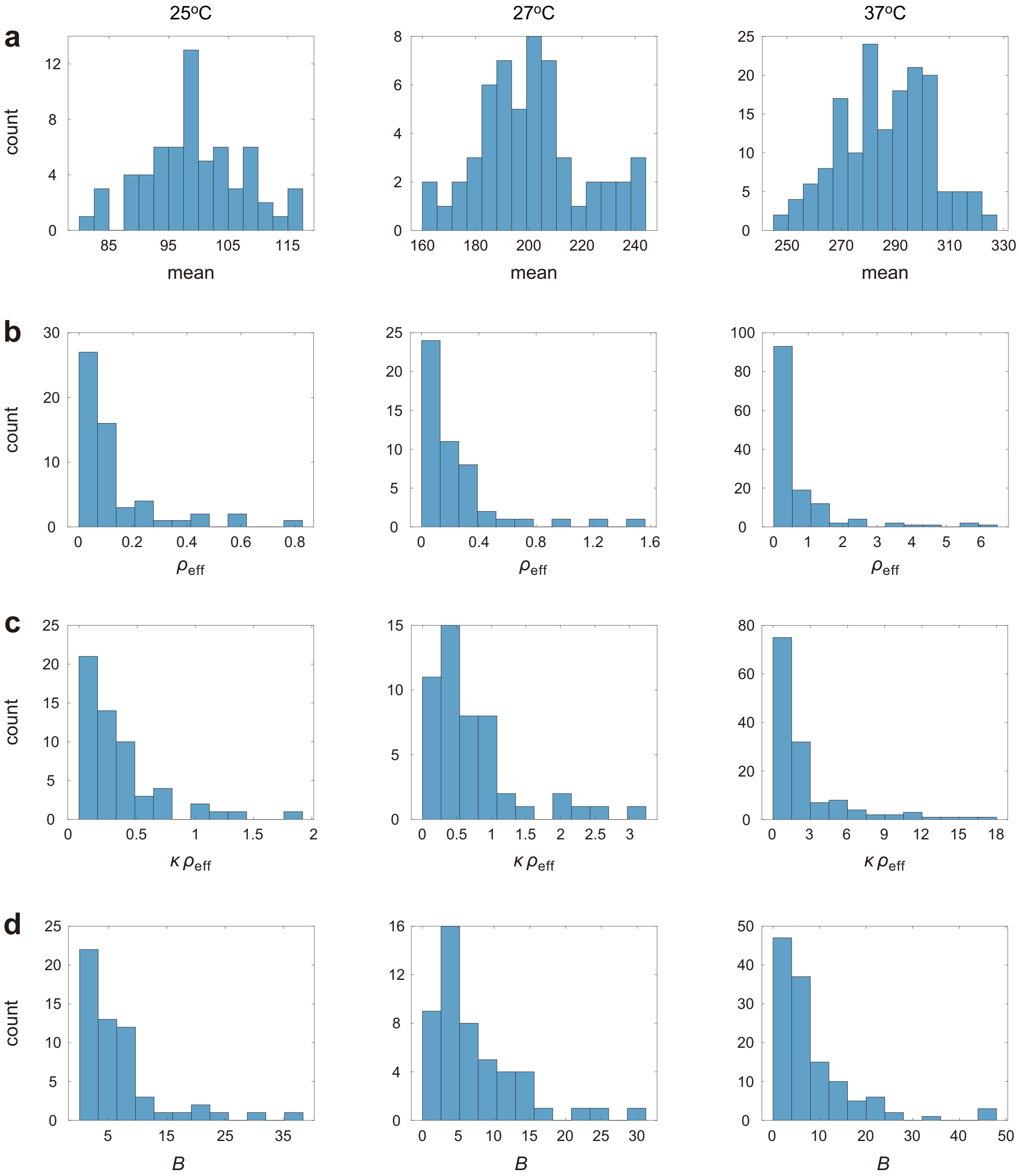


Fig. S6. **Distribution of the second set of model parameters at the three temperatures.** (a) Distribution of the estimated $\langle n \rangle$. (b) Distribution of the estimated $\rho_{\mathrm{eff}}$. (c) Distribution of the estimated $\kappa\rho_{\mathrm{eff}}$. (d) Distribution of the estimated $B$.

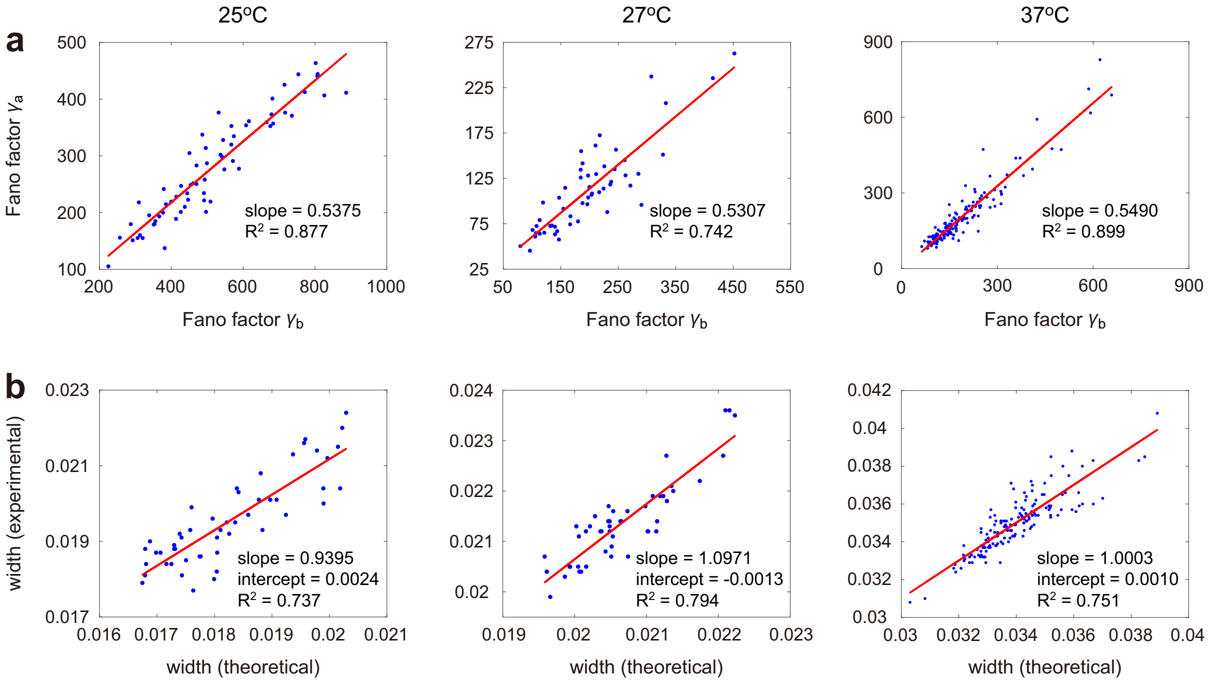


Fig. S7. **Analysis of single-cell gene expression data in *E. coli* published in [8].** (a) Comparison between the Fano factor $\gamma_b$ of fluorescence intensities just before division and the Fano factor $\gamma_a$ of fluorescence intensities just after division for all cell lineages at the three temperatures. (b) Comparison between the widths of the experimental power spectra obtained using the AR model technique and the theoretical power spectra determined using all the estimated parameters for all cell lineages at the three temperatures.
